# PAN2-PAN3 deadenylase activity is selectively required for mitotic robustness under microtubule stress

**DOI:** 10.1101/2024.05.01.591760

**Authors:** Jigyasa Verma, Zhengcheng He, Joshua A.R. Brown, Pamela Dean, Barry P. Young, Stephane Flibotte, LeAnn J. Howe, Christopher D. Maxwell, Calvin D. Roskelley, Christopher J.R. Loewen

## Abstract

**HIGHLIGHTS:**

- PAN2-PAN3 catalytic activity is required for mitotic robustness under MT stress
- Catalytic-dead Pan2-D1020A phenocopies *pan2Δ*: spindle defects and G2/M arrest
- Loss of Pan2 catalytic activity amplifies nocodazole-induced changes in *CLB1* and *CLN3* mRNA
- *PAN2* depletion in human cells produces multipolar spindles under MT stress

**IN BRIEF:** *Verma et al. show that* PAN2-PAN3 *deadenylase catalytic activity is selectively required for mitotic fidelity under microtubule stress. Yeast pan2Δ cells exhibit spindle defects, G2/M arrest, and altered CLB1 and CLN3 mRNA levels under nocodazole. PAN2 knockdown in human cells produces multipolar spindles, demonstrating evolutionary conservation*.

Cytoplasmic deadenylation by PAN2-PAN3 and CCR4-NOT regulates mRNA stability and translation, but the contribution of individual deadenylases to mitotic progression is poorly defined. Here we show that PAN2-PAN3 catalytic activity is selectively required for mitotic robustness under microtubule stress. In budding yeast, the catalytically inactive Pan2-D1020A mutant fails to complement *pan2Δ* nocodazole sensitivity, leading to G2/M arrest, spindle defects, and increased cell death. *pan2Δ* exhibits synthetic genetic interactions with tubulin and tubulin-folding genes, and RNA-seq reveals dysregulation of cell-cycle transcripts including the cyclins *CLB1* and *CLN3*. The phenotype is stress-conditional: *pan2Δ* cells proliferate normally without drug. In mammalian cells, *PAN2* depletion sensitizes HEK293A cells to nocodazole and colchicine and produces prolonged metaphase with multipolar spindles. Together, these findings define a conserved, stress-conditional role for PAN2-PAN3 deadenylase activity in safeguarding mitotic fidelity when microtubules are compromised.

Graphical Abstract:
See accompanying graphical abstract file.

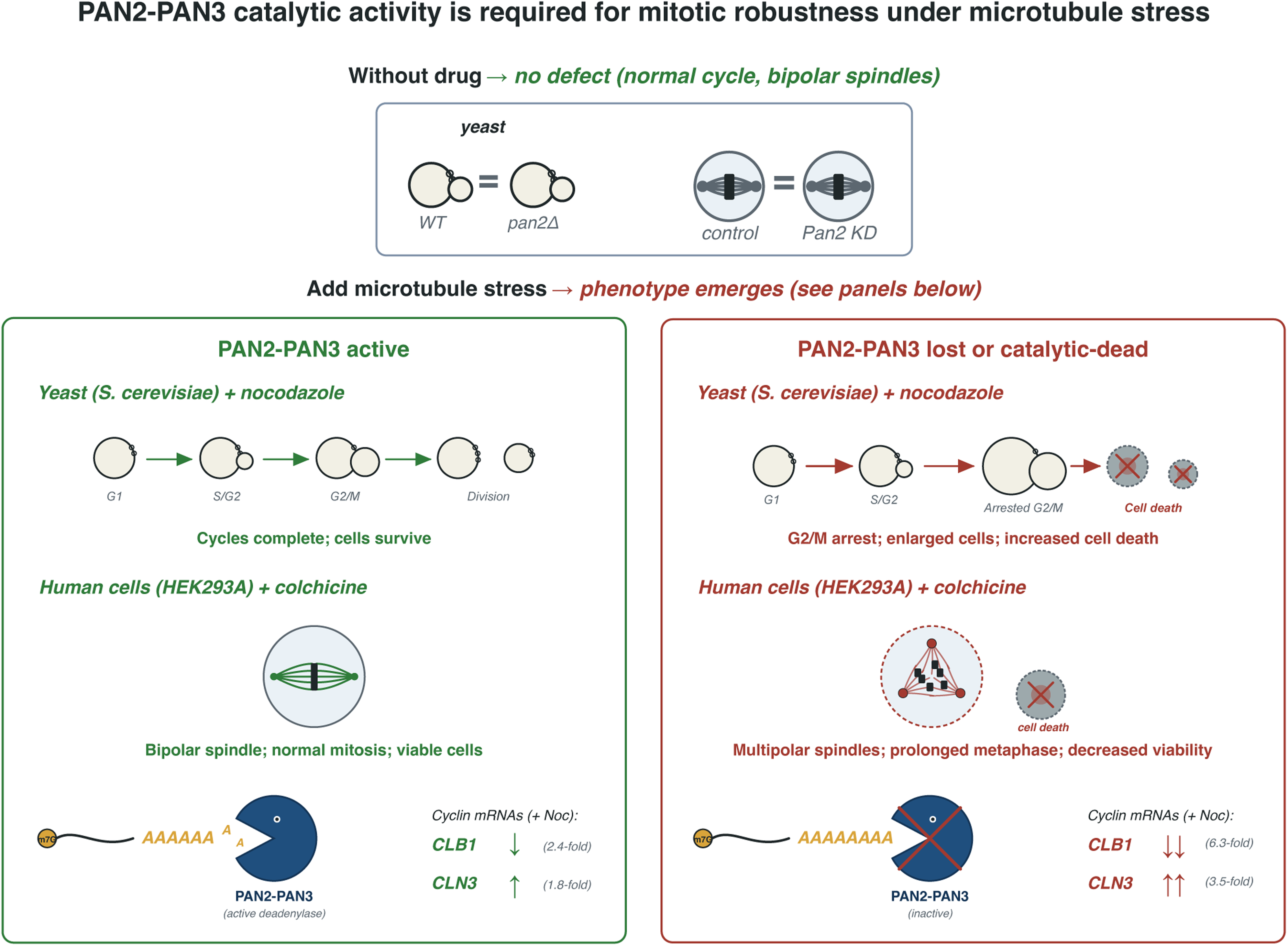

## INTRODUCTION

The poly(A) tail is a critical determinant of mRNA stability and translation, and its progressive shortening by cytoplasmic deadenylases provides a mechanism for post-transcriptional control of gene expression (Passmore and Coller, 2022). Two multi-subunit complexes, PAN2-PAN3 and CCR4-NOT, execute the bulk of cytoplasmic deadenylation, with PAN2-PAN3 trimming long poly(A) tails and CCR4-NOT carrying out subsequent shortening (Yamashita *et al*., 2005; Webster *et al*., 2018).

Post-transcriptional regulation features prominently in cell-cycle control. CCR4-NOT regulates expression of spindle assembly checkpoint components (Takahashi *et al*., 2012) and modulates cell proliferation through both deadenylation-dependent and -independent mechanisms (Aslam *et al*., 2009; Collart, 2016). Recent work has revealed that deadenylase activity is itself dynamically regulated across the cell cycle: a regulator named *MARTRE* attenuates CCR4-NOT activity during developmental transitions (Yang *et al*., 2025); global deadenylation is suppressed during mitosis to stabilize the mitotic transcriptome (Khalizeva *et al*., 2025); and distinct waves of mRNA decay occur in mitosis and early G1 (Krenning *et al*., 2022).

By contrast, the contributions of PAN2-PAN3 to cell-cycle progression remain poorly characterized. The complex comprises the catalytic exonuclease Pan2 and the regulatory pseudokinase Pan3, which together associate with poly(A)-binding proteins to engage mRNA substrates (Jonas *et al*., 2014; Schäfer *et al*., 2014; Wolf *et al*., 2014; Schäfer *et al*., 2019).

PAN2-PAN3 trims long poly(A) tails on subsets of mRNAs but exerts modest effects on global mRNA stability (Brown *et al*., 1996; Yamashita *et al*., 2005), and recent work shows that its substrate specificity is conferred by association with sequence-specific RNA-binding proteins (Tang *et al*., 2025). Genome-wide drug-sensitivity screens have placed yeast *pan2Δ* and *pan3Δ* among the most nocodazole- and benomyl-sensitive deletion strains (Hillenmeyer *et al*., 2008; Lee *et al*., 2014), and *PAN2* exhibits genetic interactions with tubulin and tubulin-folding genes (Costanzo *et al*., 2016). Whether these observations reflect a functional role in mitotic progression has not been examined.

Here, we use chemical-genetic, cell biological, and transcriptomic approaches to establish that PAN2-PAN3 deadenylase catalytic activity is selectively required for mitotic robustness under microtubule stress in budding yeast, and that *PAN2* depletion confers analogous mitotic vulnerability in human cells.

## RESULTS

### *PAN2-PAN3* deadenylase activity is required for growth under microtubule stress

A large-scale chemical-genetic screen of ∼4,800 yeast deletion mutants revealed that homozygous deletion of *PAN2* or *PAN3* ranks among the top 20 most nocodazole- and benomyl-sensitive strains, placing them within the ‘tubulin folding and SWR complex’ response signature (Hillenmeyer *et al*., 2008; Lee *et al*., 2014) (Figure S1A). Top-ranked strains also included deletions of *PAC2* (encoding an α-tubulin-binding protein), *CIN1* (β-tubulin-binding protein), *CIN2* (GTPase activator of tubulin complex assembly), and *TUB3* (α-tubulin itself) (Hoyt *et al*., 1990; Hoyt *et al*., 1997; Tian *et al*., 1996; Siegers *et al*., 1999), suggesting a role for the PAN complex in microtubule-related functions.

The screen was performed at 6 μM nocodazole, far below the 40 μM typically used to induce G2/M arrest (Amberg *et al*., 2006). To recapitulate screen conditions, log-phase *pan2Δ* and *pan3Δ* cells were treated with 6 μM nocodazole. Both mutants grew substantially slower than wild-type in drug but were indistinguishable from wild-type in DMSO vehicle (Figure 1A). A ∼100-minute lag preceded the onset of the drug effect following addition, precluding determination of exponential growth constants.

**Figure 1:**
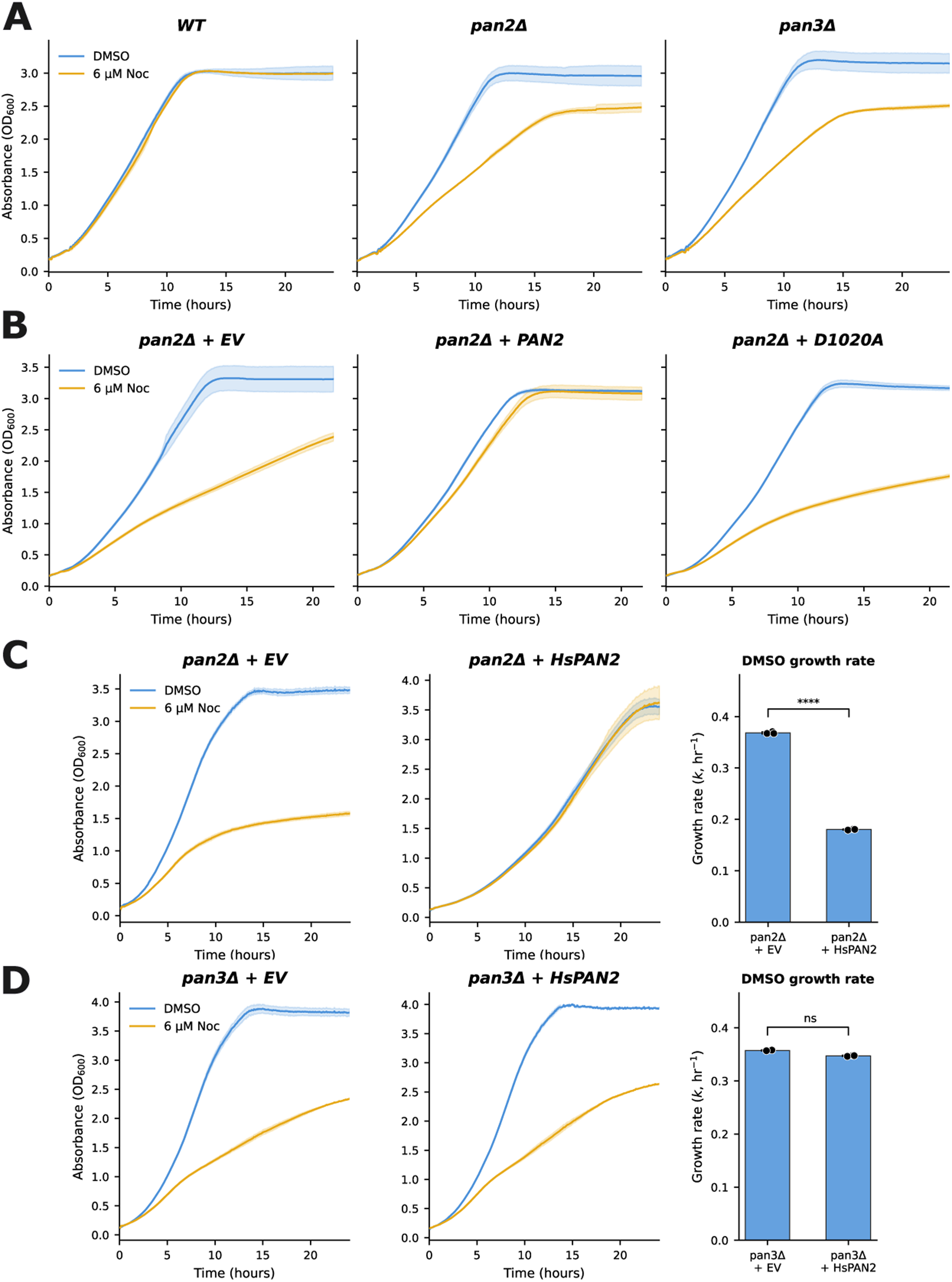
PAN2-PAN3 deadenylase activity is required for growth under microtubule stress. **(A)** Growth curves of wild-type, *pan2Δ*, and *pan3Δ* cells in SC media + 6 μM nocodazole (Noc; orange) or DMSO control (blue) at 30 °C over 24 hours. Wild-type cells exhibit comparable growth in both conditions; *pan2Δ* and *pan3Δ* cells grow substantially slower in nocodazole, with a ∼100-min lag preceding the onset of the drug effect. Absorbance (OD₆₀₀) plotted with shaded ± SD. Data: 6 technical replicates per experiment, 3 biological replicates. **(B)** Complementation: growth curves of *pan2Δ* cells expressing empty vector, wild-type Pan2, or catalytically inactive Pan2-D1020A from a centromeric (CEN) plasmid, cultured with 6 μM nocodazole or DMSO. Wild-type Pan2 restores nocodazole resistance; Pan2-D1020A fails to rescue. The Pan2-D1020A mutant carries a single amino-acid substitution within the active-site DEDD motif that abolishes deadenylation of poly(A) RNA *in vitro*. GFP-tagged variants of both proteins are expressed at comparable levels (Figure S1B). Absorbance plotted with shaded ± SD. Data: 6 technical replicates per experiment, 3 biological replicates. **(C)** Functional conservation. Left and middle: growth curves of *pan2Δ* cells expressing empty vector or codon-optimized human *PAN2* (*HsPAN2*) from a high-copy 2μ plasmid with 6 μM nocodazole or DMSO. Right: DMSO growth-rate (k, h⁻¹) comparison. *HsPAN2* rescues nocodazole sensitivity comparably to yeast Pan2 but also attenuates basal growth in DMSO, indicating dosage-dependent effects. An analogous slow-growth phenotype is observed upon overexpression of yeast *PAN2* (Figure 6D). Mean ± SD; ****p < 0.0001 by two-tailed Student’s t-test. Data: 6 technical replicates per experiment, 3 biological replicates. **(D)** Pan3 requirement. Left and middle: growth curves of *pan3Δ* cells expressing empty vector or codon-optimized human *PAN2* (*HsPAN2*) from a 2μ plasmid with 6 μM nocodazole or DMSO. Right: DMSO growth-rate (k, h⁻¹) comparison. *HsPAN2* expression in a *pan3Δ* background neither rescues nocodazole sensitivity nor produces the marked basal growth reduction seen in *pan2Δ* + *HsPAN2* (Figure 1C); growth rates of *pan3Δ* + EV and *pan3Δ* + *HsPAN2* in DMSO are indistinguishable, indicating that both phenotypes require an intact PAN2-PAN3 complex. Mean ± SD; n.s., not significant. Data: 6 technical replicates per experiment, 3 biological replicates. CEN, centromeric; EV, empty vector; *HsPAN2*, human *PAN2*; SC, Synthetic Complete media.

To test whether the phenotype depends on deadenylase catalytic activity, *pan2Δ* cells were complemented with CEN plasmids expressing wild-type Pan2 or a catalytically inactive *DEDD*-motif mutant (Pan2-D1020A) previously shown to be unable to deadenylate poly(A) RNA *in vitro* (Schäfer *et al*., 2019; Boeck *et al*., 1996). Wild-type Pan2 fully restored nocodazole resistance, whereas Pan2-D1020A did not (Figure 1B). GFP-tagged versions of both proteins were expressed at equivalent levels (Figure S1B); GFP-Pan2, but not GFP-Pan2(D1020A), rescued nocodazole sensitivity (Figure S1C), confirming that rescue depends on catalytic activity rather than protein abundance.

To assess evolutionary conservation, we expressed a yeast-codon-optimized human *PAN2* (*HsPAN2*; sequence in SFile 1) in *pan2Δ* cells from a high-copy 2μ plasmid. *HsPAN2* rescued nocodazole sensitivity to a level comparable to yeast Pan2 but also attenuated basal growth in the absence of drug (Figure 1C). This slow-growth phenotype was Pan3-dependent: *HsPAN2* expression in *pan3Δ* cells neither slowed growth nor rescued nocodazole sensitivity (Figure 1D), indicating the effect requires an intact PAN2–PAN3 complex. An analogous slow-growth phenotype was observed upon overexpression of yeast Pan2 (see Figure 6D), consistent with PAN dosage modulating proliferative capacity.

### PAN2 and PAN3 genetically interact with tubulin and genes governing tubulin folding

To understand how PAN function relates to microtubule homeostasis, we examined genetic interactions of *PAN2* and *PAN3* reported in CellMap (Costanzo *et al*., 2016) (Figure S2A). Gene-ontology analysis implicated PAN in chromosome segregation (*APC1, BUB3, CBF1, MPS1, SLI15, SMC3, TUB2, TUB3*), cytoskeletal protein binding (*ARP2, CIN1, GIM3, GIM5, PAC10, RBL2, YKE2*), secretion (*ACT1, SEC1, SSO2, VPS27*), and RNA polymerase-II-specific transcription factor binding (*CBF1, GCR2, SWI1*) (Figure S2A). Within the chromosome-segregation network, subcategories included spindle, microtubule cytoskeleton, and mitotic spindle assembly checkpoint signaling, consistent with a role for PAN in spindle function.

To validate these genetic interactions and extend them to the microtubule-stress regime, we performed synthetic genetic array (SGA) screens of a *pan2Δ* query against the non-essential deletion collection (∼4,800 strains) in the absence and presence of a low dose of nocodazole (1.5 μM) that does not produce a growth defect in *pan2Δ* alone (Figure S2B). In the absence of nocodazole, using an aggravating cutoff of 0.7 (three standard deviations; 99.7% CI) and alleviating cutoff of 1.3 at p < 0.05 across 3/3 replicates, we identified 52 aggravating and 27 alleviating interactors (SFile 2); four (*TUB3, BUB3, RPL12B, RPL14A*) overlapped with the previously reported *PAN2* interactome (Costanzo *et al*., 2016) (Figure 2A). Notably, most chromosome-segregation interactors were not recovered without nocodazole.

**Figure 2:**
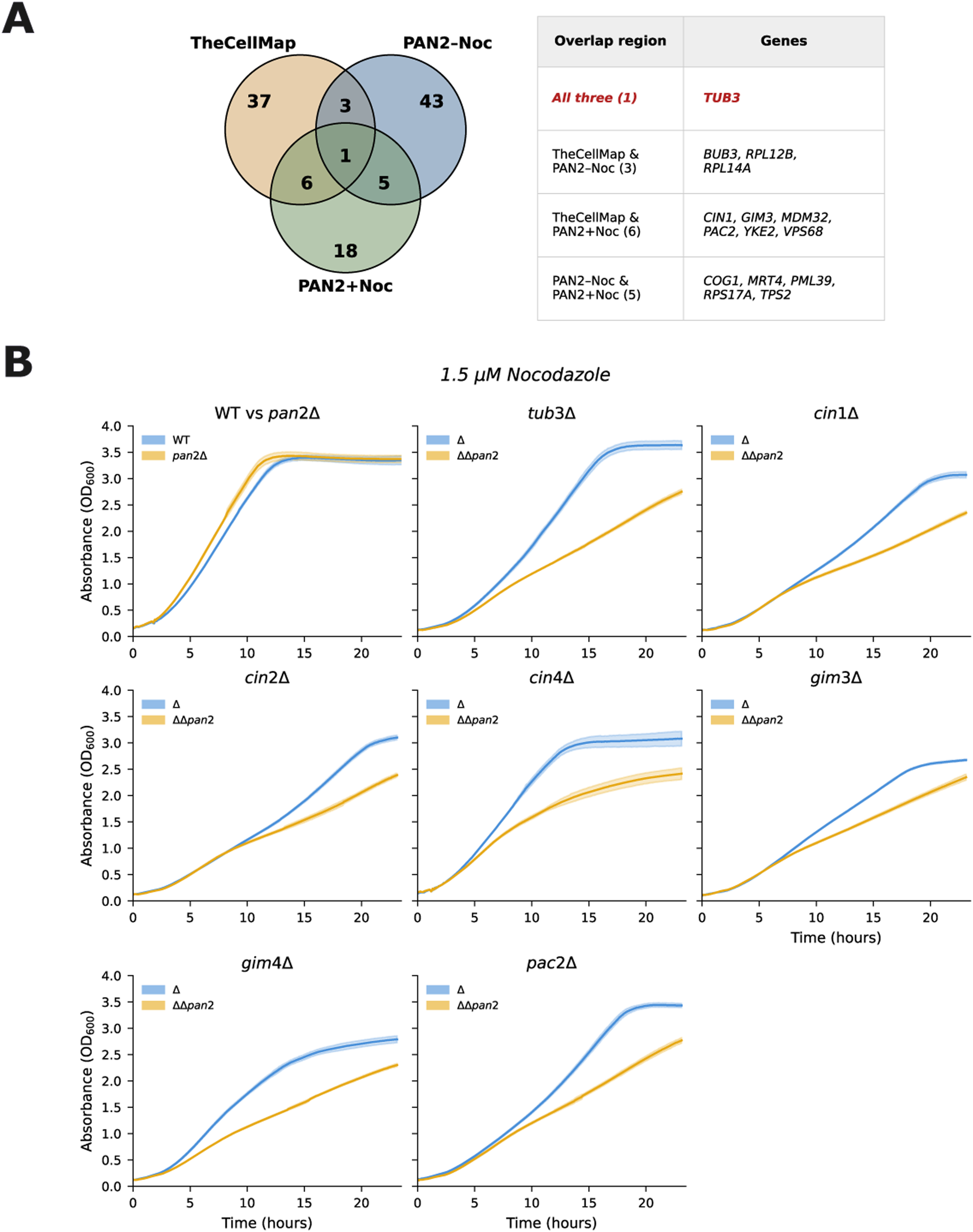
*PAN2* and *PAN3* genetically interact with tubulin and tubulin-folding genes. **(A)** Venn diagram of *pan2Δ* genetic interactions identified by synthetic genetic array (SGA) screens against the non-essential deletion collection (∼4,800 strains) under DMSO and 1.5 μM nocodazole conditions, intersected with previously reported *PAN2* interactions in CellMap (Costanzo *et al*., 2016). Aggravating cutoff 0.7 (three SD; 99.7% CI) and alleviating cutoff 1.3 at p < 0.05 across 3/3 replicates yielded 52 aggravating + 27 alleviating interactors in DMSO and 30 aggravating + 21 alleviating in 1.5 μM nocodazole. The right table summarizes overlap genes by region: 1 shared across all three datasets (*TUB3*); 3 shared between DMSO SGA and CellMap (*BUB3*, *RPL12B*, *RPL14A*); 6 shared between Noc SGA and CellMap (*CIN1*, *GIM3*, *MDM32*, *PAC2*, *YKE2*, *VPS68*); and 5 between Noc and DMSO SGA only (*COG1*, *MRT4*, *PML39*, *RPS17A*, *TPS2*). **(B)** Double-mutant growth curves at 1.5 μM nocodazole. Eight panels comparing single-mutant (blue) and *pan2Δ* double-mutant (orange) growth in WT vs *pan2Δ*, *tub3Δ*, *cin1Δ*, *cin2Δ*, *cin4Δ*, *gim3Δ*, *gim4Δ*, and *pac2Δ* backgrounds, all in SC media + 1.5 μM nocodazole at 30 °C. Double mutants exhibit aggravated growth defects relative to either single mutant alone, validating the SGA hits and the expanded test set (*CIN2*, *CIN4*, *GIM4*) for PAN-tubulin-folding genetic interactions. DMSO controls for the same double mutants are shown in Figure S2B. Absorbance (OD₆₀₀) plotted with shaded ± SD. Data: 6 technical replicates per experiment, 3 biological replicates.

In the presence of 1.5 μM nocodazole, the same cutoffs yielded 30 aggravating and 21 alleviating interactors (SFile 2), of which 7 overlapped with CellMap PAN interactions (Figure 2A): *CIN1*, *GIM3*, *MDM32*, *PAC2*, *TUB3*, *VPS68*, and *YKE2*. *CIN1* encodes tubulin-folding cofactor D, which facilitates β-tubulin folding (Tian *et al*., 1996); *GIM3*, *PAC2*, and *YKE2* encode subunits of the prefoldin co-chaperone complex involved in tubulin folding (Siegers *et al*., 1999); and *TUB3* encodes α-tubulin. The requirement for nocodazole to recover microtubule-related interactions, in contrast to earlier screens (Costanzo *et al*., 2016), likely reflects differences in query-strain background, media composition, temperature, and array density between studies.

We confirmed these interactions in liquid growth assays at 1.5 μM nocodazole: *pan2Δ* double mutants with *CIN1*, *GIM3*, *PAC2*, or *TUB3* exhibited aggravated growth defects relative to either single mutant (Figure 2B). We also extended the test set to additional tubulin-folding genes, *CIN2* (GTPase-activating protein / tubulin folding cofactor C), *CIN4* (GTPase regulating β-tubulin folding under *CIN2* control), and *GIM4* (prefoldin complex subunit), which similarly displayed aggravating interactions with *PAN2* under nocodazole but not in its absence (Figure S2B). Because PAN regulates mRNA stability, we tested whether PAN loss alters tubulin-transcript abundance. qRT-PCR at 6 hours post-treatment showed no significant change in *TUB1*, *TUB2*, or *TUB3* mRNA between *pan2Δ* + Pan2-D1020A and *pan2Δ* + Pan2-WT cells under either DMSO or 6 μM nocodazole conditions (Figure S2C), ruling out catalytic-activity-dependent regulation of tubulin transcripts and implicating post-transcriptional mechanisms acting on other targets.

### PAN2 deletion exacerbates nocodazole-induced G2/M arrest

To determine the cell-cycle consequences of *PAN2* loss, we performed flow cytometry on log-phase *pan2Δ* and wild-type cells treated with DMSO or 6 μM nocodazole for 3, 6, or 10 hours. Cells were fixed in ethanol and stained to analyze DNA content. In DMSO, both strains showed the typical 1N/2N distribution of asynchronous log-phase cultures at all time points (Figure 3A). Upon nocodazole treatment, *pan2Δ* cells exhibited an earlier and more pronounced shift from 1N to 2N, with broadening of the 2N peak relative to wild-type at 3 hours, indicative of a G2/M delay (Figure 3A). By 6 and 10 hours, the 1N peak was largely lost and DNA content extended beyond 2N in *pan2Δ*, suggesting genomic instability or extensive G2/M arrest (Figure 3A).

**Figure 3:**
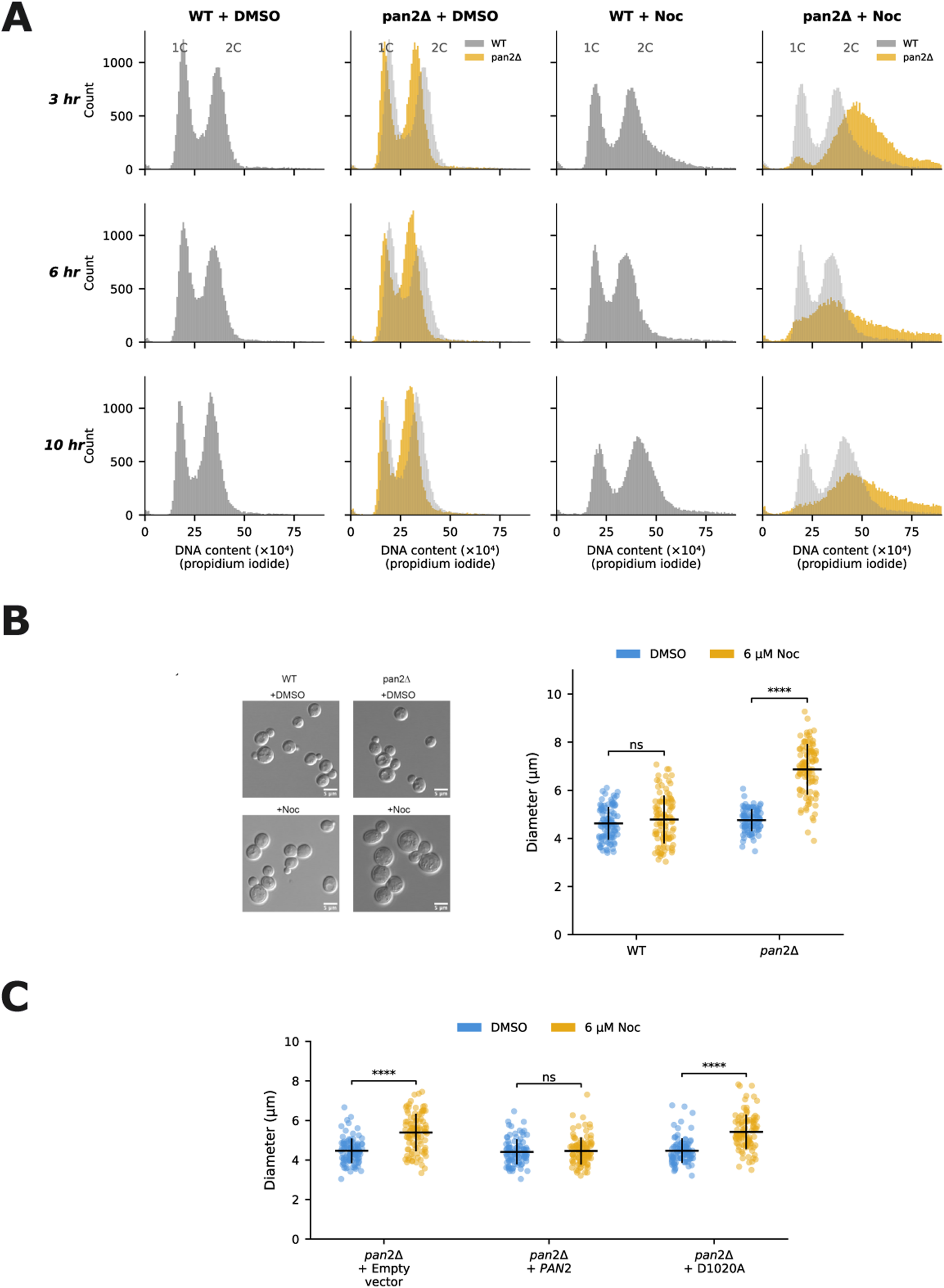
Loss of Pan2 catalytic activity causes G2/M arrest and cell enlargement. **(A)** Flow cytometry DNA content analysis. Log-phase wild-type and *pan2Δ* cells were cultured ± 6 μM nocodazole for 3, 6, and 10 hours, fixed in ethanol, and stained for DNA content (propidium iodide). Histograms show G1 (1C) and G2/M (2C) populations under four conditions (WT + DMSO, *pan2Δ* + DMSO, WT + Noc, *pan2Δ* + Noc). Wild-type cells accumulate at 2C upon nocodazole. *pan2Δ* cells show an earlier, more pronounced shift from 1C to 2C with broadening of the 2C peak at 3 hours; by 6 and 10 hours the 1C peak is largely lost and DNA content extends beyond 2C, suggesting genomic instability or extensive G2/M arrest. Gray WT overlay on *pan2Δ* columns facilitates direct comparison. Data: representative histograms from 3 biological replicates; ≥10,000 cells per sample. **(B)** Cell diameter analysis. Representative bright-field images (left) and quantification of cell diameter (right) for WT and *pan2Δ* cells cultured with 6 μM nocodazole or DMSO for 6 hours. Under nocodazole, *pan2Δ* cells exhibit a ∼1.75-fold increase in cell diameter relative to WT and are predominantly large-budded, consistent with G2/M delay. Mean ± SD; n.s., not significant; ****p < 0.0001 by two-tailed Welch’s t-test. Scale bar = 5 μm. Data: n > 100 cells per condition; 3 biological replicates. **(C)** Complementation of cell-size phenotype. Cell diameter of *pan2Δ* + empty vector, *pan2Δ* + Pan2-WT, and *pan2Δ* + Pan2-D1020A cells cultured with 6 μM nocodazole or DMSO for 6 hours. Wild-type Pan2 restores cell size toward WT levels, whereas Pan2-D1020A does not, indicating that the enlargement phenotype requires Pan2 catalytic activity. Mean ± SD; n.s., not significant; ****p < 0.0001 by two-tailed Welch’s t-test. Data: n > 100 cells per condition; 3 biological replicates.

Forward-scatter analysis during flow cytometry showed that nocodazole-treated *pan2Δ* cells had elevated forward scatter relative to wild-type as early as 3 hours after treatment, consistent with increased cell size, while DMSO-treated cells showed no size difference; this trend persisted at 6 and 10 hours (Figure S3A). Morphometric analysis at 6 hours confirmed a ∼1.75-fold increase in cell diameter in nocodazole-treated *pan2Δ* cells relative to wild-type (Figure 3B); these cells were predominantly large-budded, consistent with G2/M delay. Complementation with wild-type Pan2, but not Pan2-D1020A, restored cell size toward wild-type levels (Figure 3C), demonstrating that the enlargement phenotype, like the arrest itself, requires Pan2 catalytic activity.

### Loss of Pan2 leads to defective mitotic progression and increased cell death under microtubule stress

To examine spindle morphology in *pan2Δ* cells under nocodazole stress, we tagged tubulin by integrating an N-terminal yomWasabi-Tub1 fusion (with native 3′UTR) (Markus *et al*., 2015). The *pan2Δ*-tagged strain retained nocodazole sensitivity comparable to untagged *pan2Δ*, while the WT-tagged strain showed only a mild basal slow-growth phenotype (Figure S4A). Live-cell imaging of these strains, with quantification by a semi-automated image-analysis pipeline (see Methods), was used to assess total cell number, dead cell counts, and cell area. In the absence of drug, wild-type and *pan2Δ* cells showed no significant differences in cell number, death, or size over 8 hours of imaging (Figure 4A, left panels; SFiles 4 and 6). Initial imaging at 6 μM nocodazole also failed to produce significant differences between strains (Figure S4B–S4C), a discrepancy from the liquid growth assay that we attributed to the static, non-aerated imaging conditions reducing effective drug exposure. We therefore increased the dose to 12 μM.

**Figure 4:**
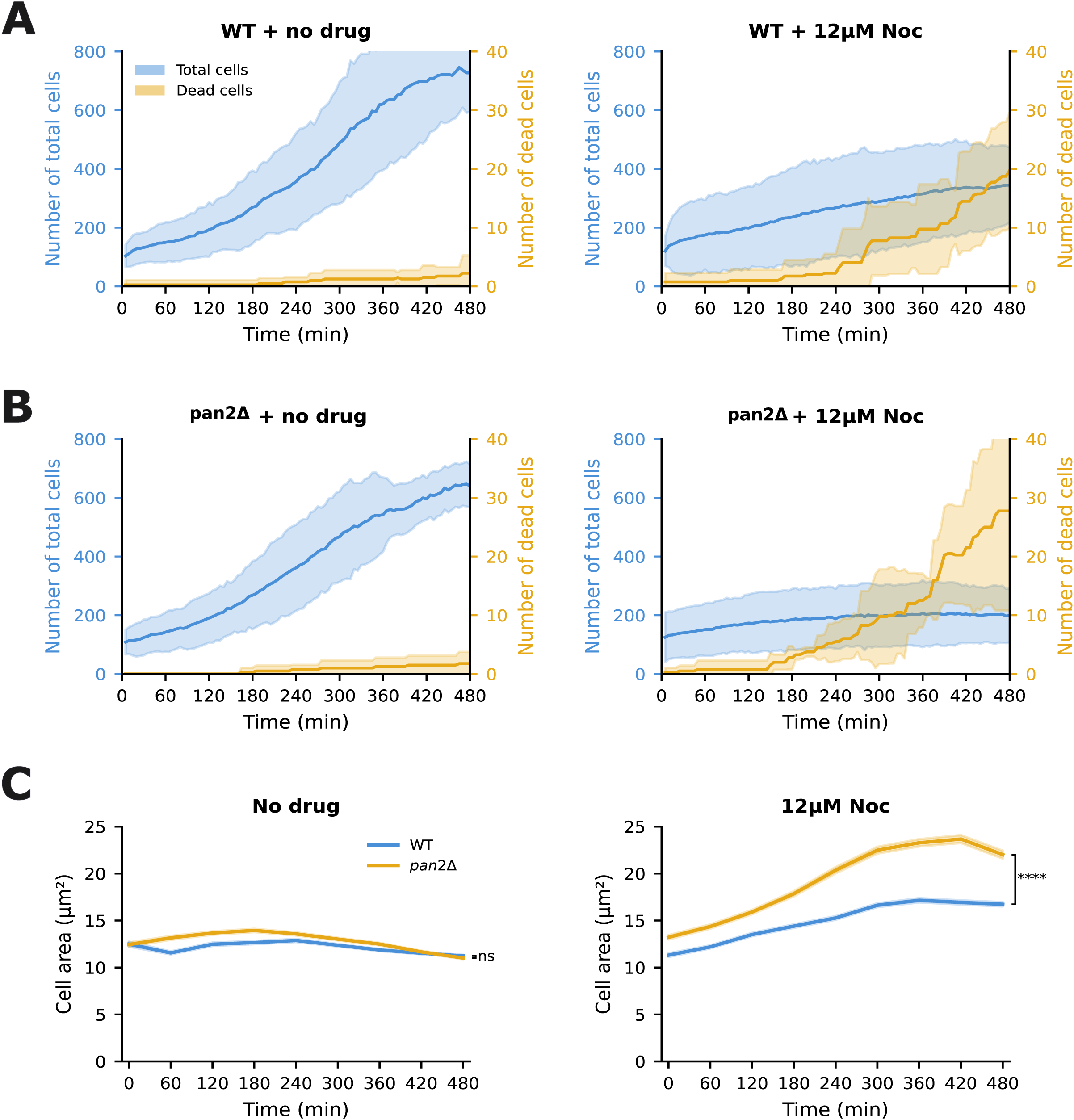
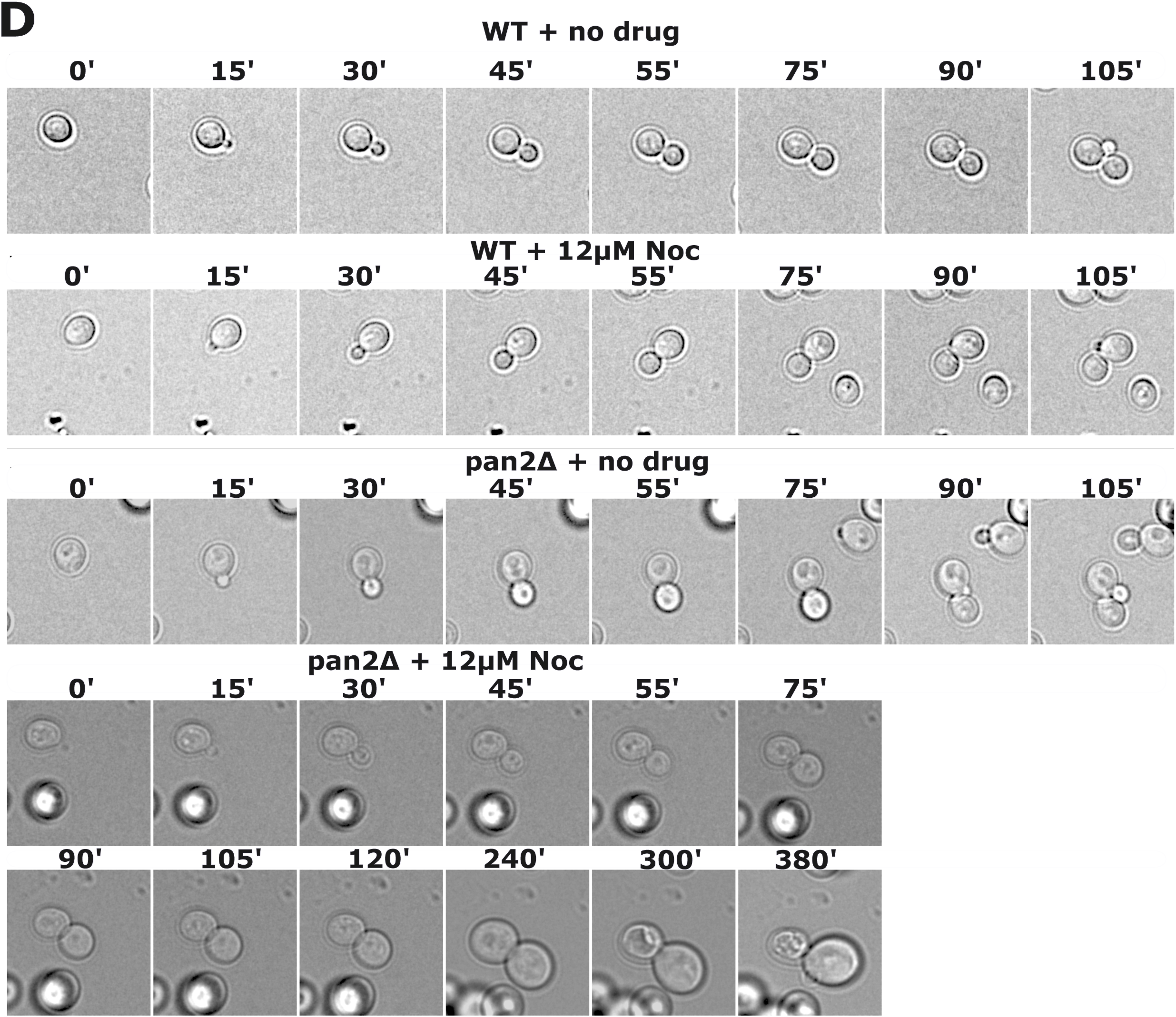
Loss of Pan2 leads to defective mitotic progression and increased cell death under microtubule stress. **(A)** Live-cell imaging quantification of WT cells. Total cell count (left y-axis, blue) and dead cell count (right y-axis, orange) over 480 minutes (8 hours) under no drug (left) or 12 μM nocodazole (right). Under 12 μM nocodazole, WT mean cell number increases from 120 (t = 0) to 344.5 (t = 480 min) while mean dead cells increase from 0.75 to 19.75. Quantification by a semi-automated image-analysis pipeline (see Methods). Mean ± 95% confidence interval (t-distribution, df = 3) shaded. Data: n = 4 biological replicates per condition. **(B)** Live-cell imaging quantification of *pan2Δ* cells, layout as in (A). Under 12 μM nocodazole, *pan2Δ* mean cell number rises only from 125 (t = 0) to 199 (t = 480 min) while mean dead cells increase from 0.25 to 27.75. Both total cell number (p < 0.0001) and dead-cell number (p < 0.0001) differ significantly from WT (Welch’s t-test on endpoint values). Mean ± 95% CI shaded. Data: n = 4 biological replicates per condition. **(C)** Mean cell area over time. Both strains increase in area with 12 μM nocodazole; *pan2Δ* cells reach significantly larger areas, plateauing at ∼360 minutes. *pan2Δ* mean cell area increases from ∼13 μm² (t = 0) to ∼23 μm² (t = 480 min); WT mean cell area increases from ∼12 μm² to ∼17 μm² over the same interval. n.s., not significant; ****p < 0.0001 by Welch’s t-test on endpoint values. Mean ± SEM (per-timepoint cell-pool SEM, SD/√n) shaded. Data: n = 380–2,726 individual cells per timepoint, 3 biological replicates. **(D)** Representative bright-field time-lapse montage of WT and *pan2Δ* cells imaged ± 12 μM nocodazole. Cells were pre-treated with DMSO or 12 μM nocodazole for ∼75 min in the imaging 96-well plate (approximately one WT cell-cycle interval) before imaging began (t = 0 in the montage). In the absence of drug, both genotypes complete a full cell-cycle within ∼75 min, with bud emergence at ∼15 min, bud growth through G2 (∼30 min), and mother–daughter separation by ∼55 min. Treated WT cells show a modestly elongated cell-cycle (∼90 min) but retain normal cell morphology and complete cytokinesis. In contrast, the representative *pan2Δ* cell under nocodazole fails to complete cell division: bud growth proceeds for ∼90 min but mother and bud continue to enlarge without separating (270 min); by 300 min the mother dies and the orphan daughter fails to grow or divide through 380 min. Time stamps in minutes (top left of each frame). See SFiles 4–7 for full movies. Scale bar = 5 μm.

At 12 μM nocodazole, wild-type and *pan2Δ* cultures differed significantly in both total cell number (p < 0.0001) and number of dead cells (p < 0.0001). Wild-type mean cell number rose from 120 at t=0 to 344.5 at t=480 min, while mean dead cells increased from 0.75 to 19.75 (Figure 4A; SFile 5). In contrast, *pan2Δ* mean cell number rose only from 125 to 199 while dead cells increased from 0.25 to 27.75 over the same interval (Figure 4B; SFile 7). Mean cell area rose in both strains but more steeply in *pan2Δ* (∼13 μm² to ∼23 μm²) than in wild-type (∼12 μm² to ∼17 μm²), plateauing at ∼360 min (Figure 4C). Together, the decreased growth, elevated death, and enlargement of nocodazole-treated *pan2Δ* cells are consistent with a defective cell cycle.

Time-lapse imaging of representative single cells corroborated these findings (Figure 4D). In untreated cells, wild-type and *pan2Δ* progressed through the cell cycle with comparable kinetics (∼75 min). Following 75 minutes of 12 μM nocodazole treatment, approximately one cell-cycle and prior to extensive cell death, wild-type cells showed a modestly elongated cycle (∼90 min) but completed mitosis with normal morphology, whereas *pan2Δ* cells frequently failed to complete division, with continued mother and bud growth followed by cell death. Together, these observations indicate that *pan2Δ* cells fail to complete mitosis under microtubule stress.

### Quantitative population-level analysis confirms disorganized microtubule structures

To determine whether the mitotic failure in *pan2Δ* cells reflects compromised microtubule organization, we performed endpoint fluorescence imaging of *TUB1*-yomWasabi cells. Unlike the live-cell imaging in Figure 4, which required 12 μM nocodazole because static, non-aerated imaging conditions reduced effective drug exposure, here cells were grown in shaking liquid culture at 6 μM nocodazole, matching the dose used in all our other yeast experiments. Cells were collected after 10 hours of treatment for endpoint imaging (dead cells excluded from analysis).

Wild-type cells retained organized microtubule structures even after prolonged nocodazole exposure, whereas *pan2Δ* cells exhibited diffuse, disorganized tubulin distribution (Figure 5A). GFP fluorescence-intensity distributions were quantified across segmented cells as a proxy for microtubule organization. Peak-to-mean intensity ratios, which capture the prominence of bright tubulin puncta relative to diffuse cytoplasmic signal, were significantly reduced in *pan2Δ* cells under nocodazole (Figure 5B), consistent with loss of discrete microtubule structures; the underlying population distribution showed *pan2Δ* + nocodazole cells uniformly clustered at low values, whereas wild-type cells retained a broad distribution with substantial fractions at high values (Figure S5A). The coefficient of variation (CV) of GFP intensity, a measure of tubulin distribution heterogeneity, was likewise reduced in *pan2Δ* + nocodazole cells (Figure 5C), indicating more uniform fluorescence, with the same population-level signature: a narrow, low-CV distribution for *pan2Δ* versus a broader, more heterogeneous distribution for wild-type (Figure S5B). Cell area measurements corroborated the enlargement phenotype (Figure 5D), with *pan2Δ* + nocodazole cells distributed toward larger sizes than wild-type (Figure S5C). Together, these population-level endpoint measurements provide independent quantitative support for a role of Pan2 in maintaining organized microtubule structures under prolonged microtubule stress.

**Figure 5:**
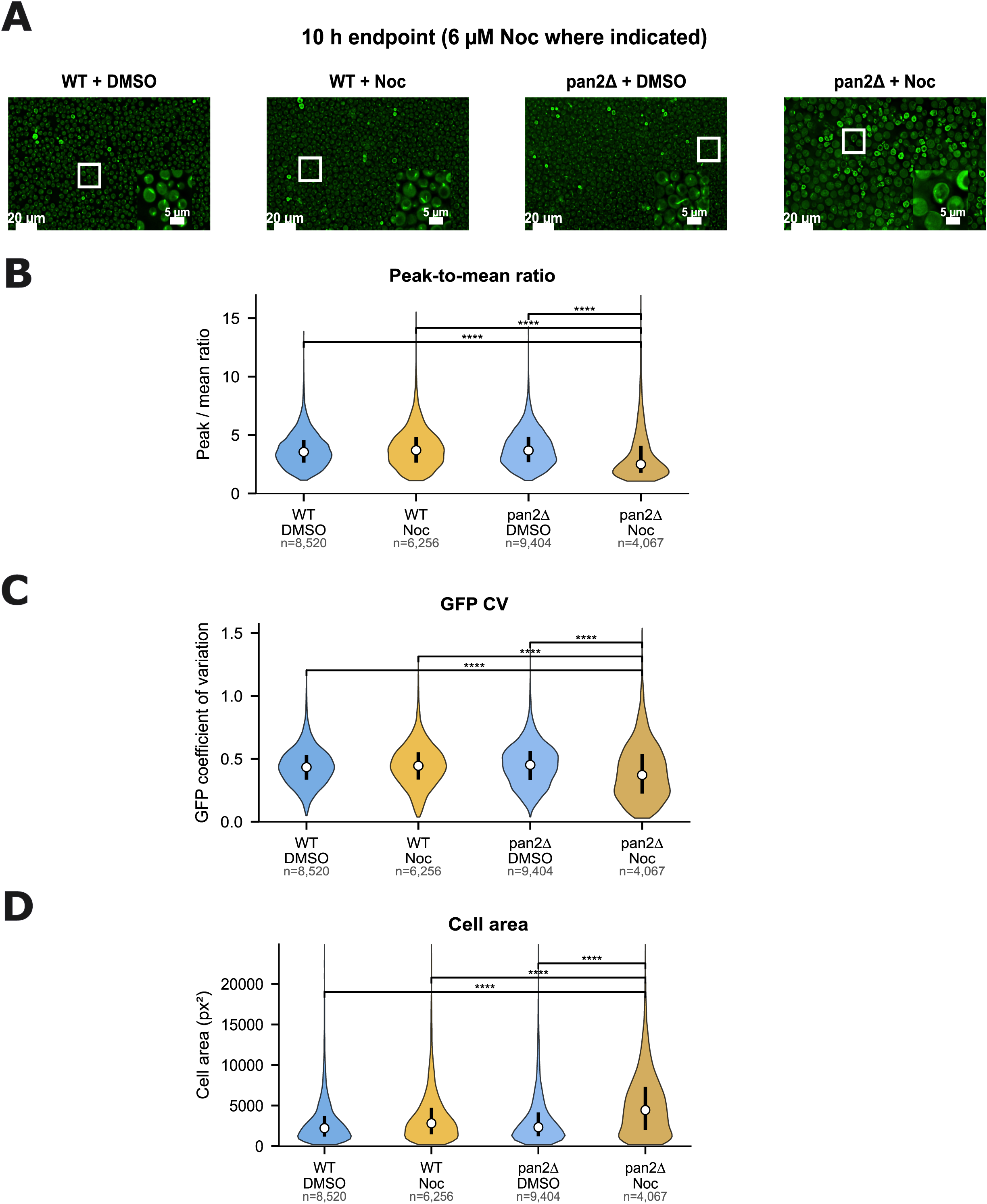
Quantitative endpoint analysis confirms impaired microtubule organization. **(A)** Representative GFP-tubulin fluorescence images of wild-type and *pan2Δ* cells expressing *TUB1*-yomWasabi after 10 hours of 6 μM nocodazole or DMSO vehicle. Wild-type cells maintain organized microtubule structures with bright puncta (spindle pole bodies and microtubule bundles) after prolonged nocodazole exposure, whereas *pan2Δ* cells exhibit diffuse, disorganized tubulin signal. Scale bar = 20 μm. **(B)** Peak-to-mean GFP intensity ratio (maximum pixel intensity / mean pixel intensity within each segmented cell) for living cells across four conditions (WT + DMSO, n = 8,520; WT + Noc, n = 6,256; *pan2Δ* + DMSO, n = 9,404; *pan2Δ* + Noc, n = 4,067), shown as violin plots. Values near 1 indicate spatially uniform fluorescence; higher values indicate concentrated signal at discrete foci. *pan2Δ* + Noc cells show significantly reduced ratios, consistent with loss of bright GFP-tubulin puncta. **(C)** Coefficient of variation (CV) of GFP intensity (standard deviation / mean across all pixels in each segmented cell), shown as violin plots; n’s as in (B). Reduced CV in *pan2Δ* + Noc cells indicates more uniform fluorescence distribution, consistent with loss of discrete microtubule structures. **(D)** Cell area (px²) from segmented masks, shown as violin plots; n’s as in (B). Increased area in *pan2Δ* + Noc cells is consistent with prolonged G2/M arrest. For panels B–D, three planned contrasts (two-tailed Welch’s t-test, Bonferroni-corrected): *p < 0.05, **p < 0.01, ***p < 0.001, ****p < 0.0001. Data pooled from 3 independent biological replicates. See also Figure S5.

### pan2Δ cells exhibit dysregulated cell-cycle gene expression in response to nocodazole

Having established that *pan2Δ* cells fail mitosis with disorganized microtubules under nocodazole, we asked what molecular changes accompany loss of Pan2 catalytic activity. We performed RNA-sequencing on duplicate log-phase cultures of *pan2Δ* + WT-Pan2 and *pan2Δ* + D1020A-Pan2 cells treated with 6 μM nocodazole for 6 hours. Spearman correlations between replicates confirmed reproducibility (Figure S6A). Of 6,031 ORFs analyzed, 129 genes were upregulated and 39 downregulated in D1020A-Pan2 cells relative to WT-Pan2 cells, using thresholds of |z-score| > 2 (equivalent to log2 fold-change cutoffs of 1.182 / −1.0554), p < 0.01, and log2 CPM > 1 (Figure 6A; SFile 3). GO enrichment of differentially expressed genes implicated replication fork protection complex, arginine biosynthetic process, carboxylic acid metabolic process, ascospore wall assembly, thiamine biosynthetic process, hexitol metabolic process, and glyoxalase III activity (Figure 6B). Replication-fork-complex transcripts, notably histone mRNAs, which are synthesized exclusively in S phase (Marzluff *et al*., 2008), are consistent with nocodazole-treated PAN-deficient cells being arrested in G2/M. Upregulation of hexitol metabolism and glyoxalase III activity points to a metabolic shift and oxidative stress response, while the increase in ascospore-wall-assembly transcripts in these haploid cells likely reflects elevated cell-wall synthesis demand accompanying sustained bud growth in arrested, enlarged cells.

**Figure 6:**
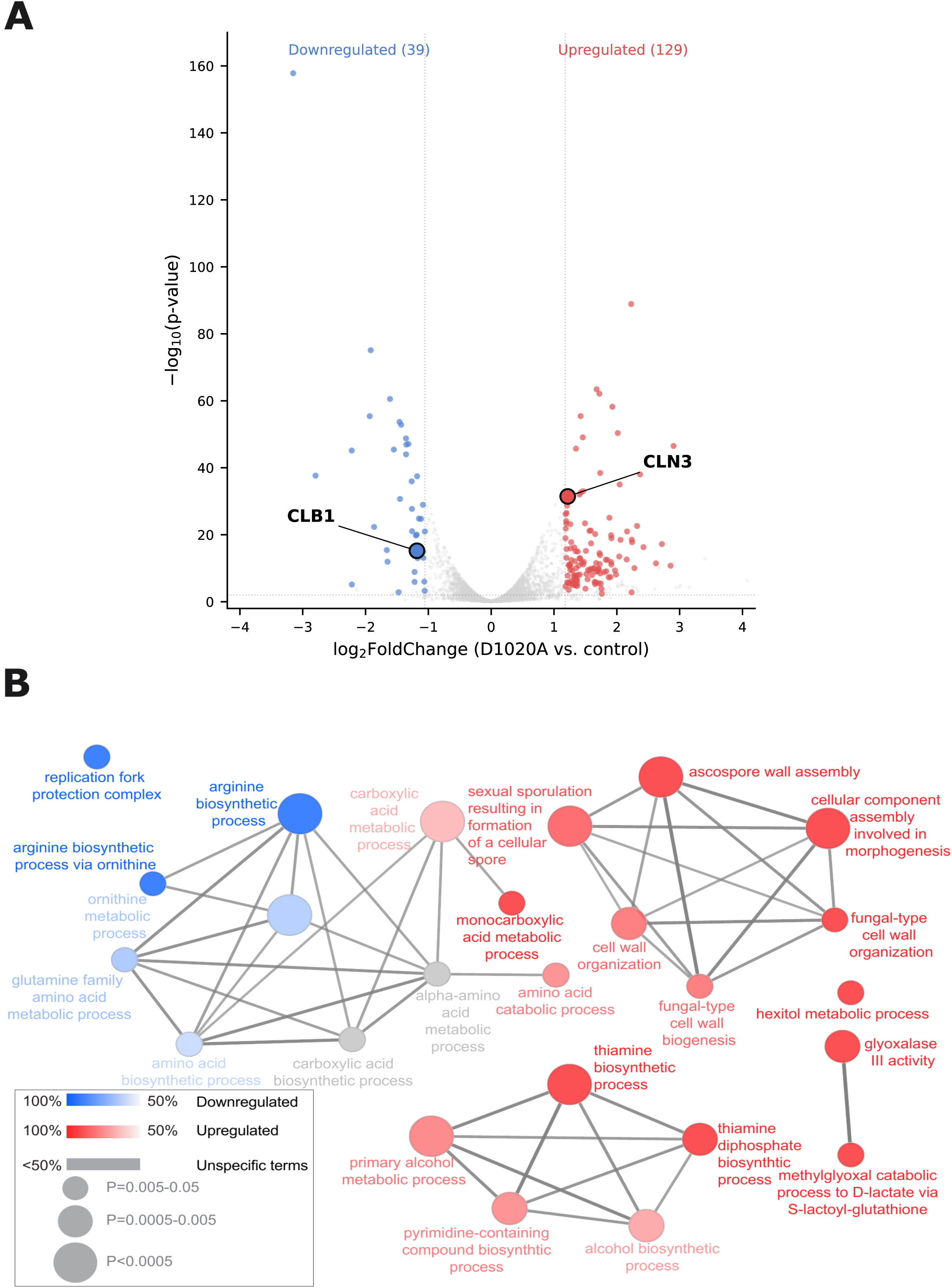
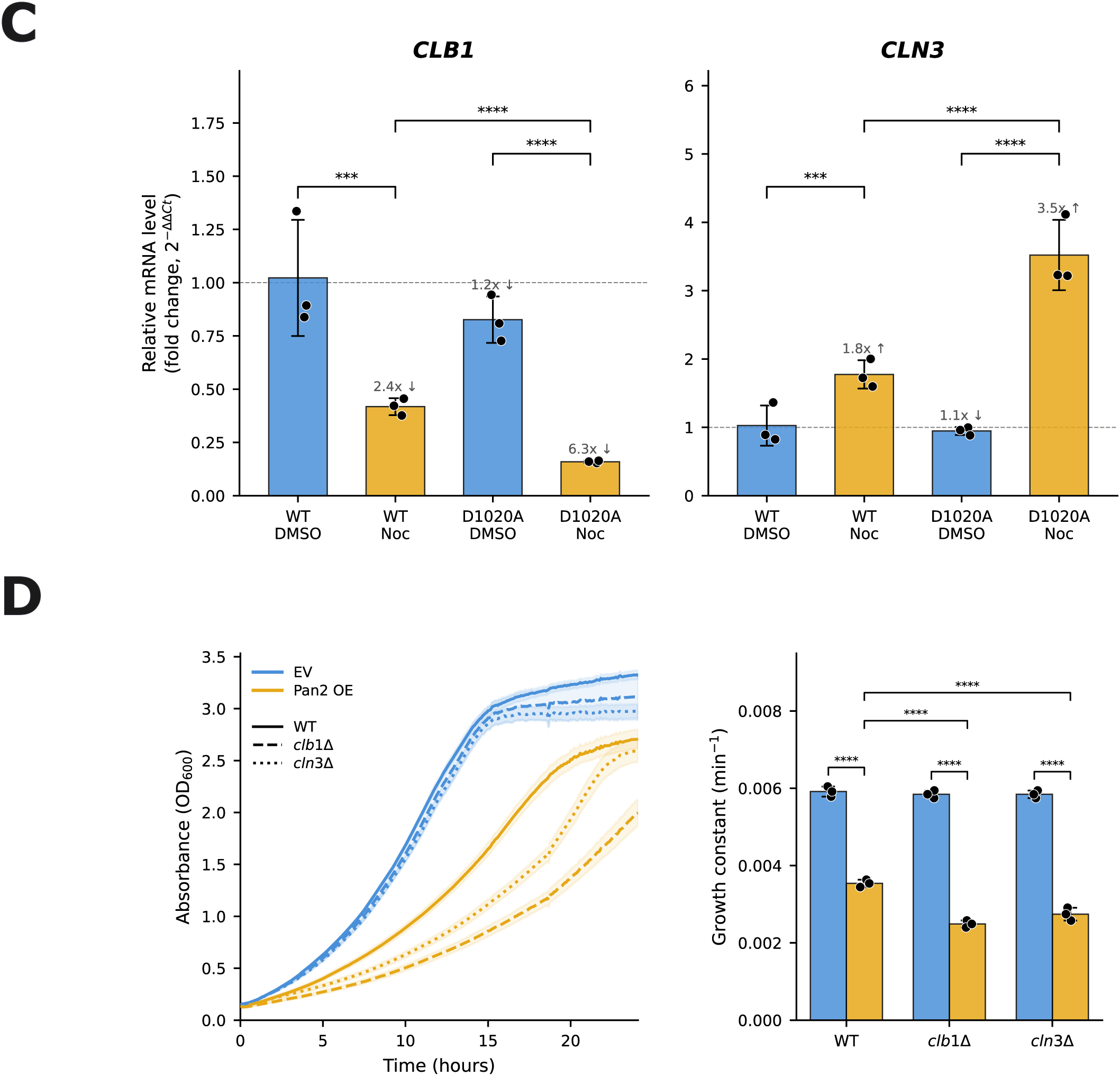
Loss of Pan2 catalytic activity is associated with altered cell-cycle gene expression under microtubule stress. **(A)** RNA-sequencing volcano plot comparing *pan2Δ* cells expressing Pan2-D1020A (catalytically inactive) versus WT Pan2, cultured with 6 μM nocodazole for 6 hours. Each point represents one of 6,031 ORFs analyzed, with log₂(fold-change) on the x-axis and –log₁₀(p-value) on the y-axis. Significantly differentially expressed genes are colored red (upregulated, n = 129) or blue (downregulated, n = 39) at |z-score| > 2 (equivalent to log₂ fold-change cutoffs of +1.180 / −1.055), p < 0.01, and log₂ CPM > 1. Cell-cycle-relevant hits *CLN3* (upregulated; G1/S cyclin) and *CLB1* (downregulated; M-phase cyclin) are circled and labeled. Data: 2 biological replicates. **(B)** Gene ontology enrichment network of differentially expressed genes (Biological Process). Enriched terms include replication fork protection complex, arginine biosynthetic process, carboxylic acid metabolic process, ascospore wall assembly, cellular component assembly involved in morphogenesis, fungal-type cell wall organization, thiamine biosynthetic process, hexitol metabolic process, glyoxalase III activity, methylglyoxal catabolic process, and primary alcohol metabolic process. Replication-fork-complex transcripts (notably histone mRNAs synthesized exclusively in S phase) are consistent with G2/M arrest of nocodazole-treated PAN-deficient cells; upregulation of hexitol metabolism and glyoxalase III activity points to a metabolic shift and oxidative stress response. Node color: blue = downregulated, red = upregulated; intensity: –log₁₀(adjusted p-value); node size: % of dysregulated genes; gray edges connect terms sharing genes; gray nodes = unspecific terms. **(C)** Quantitative RT-PCR validation of *CLB1* and *CLN3* mRNA levels in *pan2Δ* cells expressing Pan2-WT or Pan2-D1020A, ± 6 μM nocodazole for 6 hours. Fold-change calculated as 2^(−ΔΔCt) normalized to *ACT1*. *CLB1* mRNA decreases 2.4-fold in WT-Pan2 cells and 6.3-fold in D1020A-Pan2 cells; *CLN3* mRNA increases 1.8-fold in WT-Pan2 cells and 3.5-fold in D1020A-Pan2 cells. Mean ± SD; n.s., not significant; ***p < 0.001, ****p < 0.0001 by pairwise two-tailed Student’s t-test. Data: 3 biological replicates with technical triplicates. **(D)** Functional association of PAN with *CLB1* and *CLN3* by overexpression. Left: growth curves of WT, *clb1Δ*, and *cln3Δ* strains transformed with empty vector (blue) or yeast *PAN2* on a high-copy 2μ plasmid (Pan2 OE; orange), cultured in SC media at 30 °C. Right: normalized exponential growth constants for the same conditions; the reduction in growth constant upon Pan2 OE is significantly greater in *clb1Δ* and *cln3Δ* backgrounds than in WT, indicating an aggravating genetic interaction. Mean ± SD; ****p < 0.0001 by two-tailed Student’s t-test. Data: 6 technical replicates per experiment, 3 biological replicates. Quantitative epistasis analysis (observed vs. expected) is shown in Figure S6C. See also Figure S6 and SFile 3. EV, empty vector; OE, overexpression.

Two genes encoding regulators of cyclin-dependent kinase (CDK) activity, *CLB1* and *CLN3*, were differentially expressed between WT-Pan2 and D1020A-Pan2 cells under nocodazole. *CLN3* encodes a CDK regulatory subunit that governs passage through START by regulating transcription at the G1/S transition. *CLB1* encodes a B-type cyclin that regulates the G2/M transition of both the mitotic cell cycle and first meiotic division (Fitch *et al*., 1992) and contributes to mitotic spindle assembly and spindle pole body separation through its interaction with *CDK1* (Richardson *et al*., 1992). qRT-PCR confirmed the RNA-seq results. In response to nocodazole, *CLB1* mRNA decreased 2.4-fold in WT-Pan2 cells and 6.3-fold in D1020A-Pan2 cells; *CLN3* mRNA increased 1.8-fold in WT-Pan2 and 3.5-fold in D1020A-Pan2 (Figure 6C). Two-way ANOVA confirmed a significant genotype × drug interaction for both transcripts, with only minor basal differences between WT-Pan2 and D1020A-Pan2 in DMSO (≤1.2-fold, n.s.) but substantially larger nocodazole-induced changes in D1020A. Thus, loss of Pan2 catalytic activity amplifies the nocodazole-induced changes in both *CLB1* and *CLN3* mRNA levels.

To test functional links between these cyclin-expression changes and the *pan2Δ* nocodazole-sensitivity phenotype, we constructed *pan2Δ clb1Δ* and *pan2Δ cln3Δ* double mutants. Growth of each double mutant matched the *pan2Δ* single mutant in both the absence and presence of nocodazole (Figure S6B), indicating that loss of either cyclin alone does not modify *pan2Δ* sensitivity. We next overexpressed yeast *PAN2* from a 2μ plasmid. As with *HsPAN2* (Figure 1C), yeast *PAN2* overexpression produced a slow-growth phenotype in wild-type cells (Figure 6D); this phenotype was enhanced in *clb1Δ* and *cln3Δ* backgrounds (Figure 6D). Quantitative epistasis analysis confirmed that observed double-mutant growth fell significantly below the multiplicative-null expectation, indicating aggravating genetic interactions between *PAN2* overexpression and both *clb1Δ* and *cln3Δ* (Figure S6C). Collectively, these data support a model in which PAN contributes to fine-tuning Clb1 and Cln3 function during the response to microtubule stress.

### PAN2 depletion in mammalian cells increases sensitivity to microtubule-destabilizing drugs

To determine whether the requirement for Pan2 during microtubule stress is conserved in mammalian cells, we generated a stable HEK293A population in which *PAN2* was knocked down using a lentiviral shRNA targeting the *PAN2* ORF (see Methods). Western blotting confirmed ∼75% reduction in Pan2 protein in the knockdown (Pan2KD) population relative to control (Figure 7A). We tested sensitivity to the microtubule-destabilizing agents nocodazole and colchicine using the colorimetric MTT (3-(4,5-dimethylthiazol-2)-2,5-diphenyltetrazolium bromide) assay, which reads out metabolic activity of live cells as absorbance at 562 nm (Mosmann, 1983). Cells were seeded at 1,000 per well, drug was added 16 hours after plating, and MTT readings were acquired at 0, 24, 48, and 72 hours thereafter (Figure 7B).

**Figure 7:**
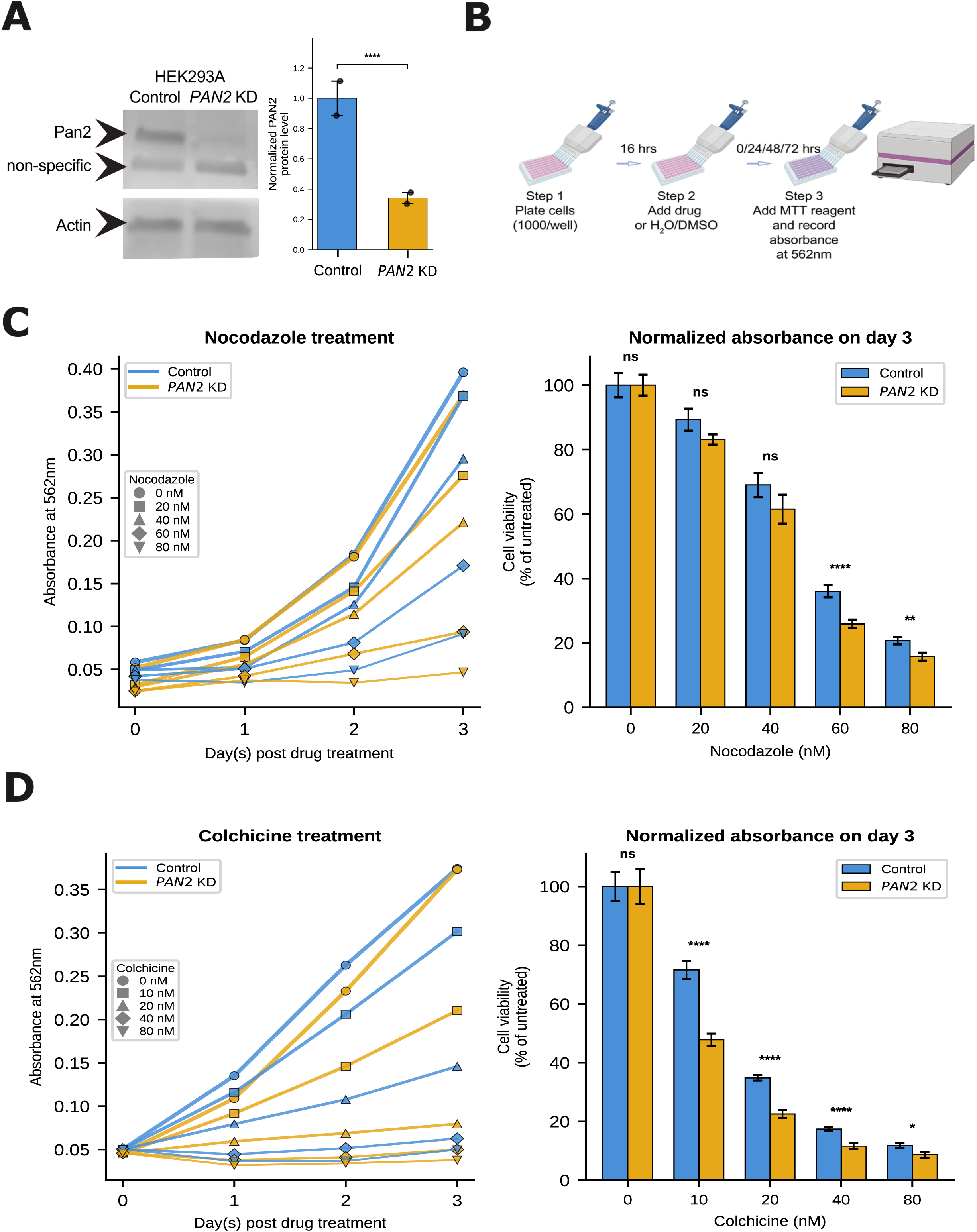
*PAN2* depletion sensitizes human cells to microtubule-destabilizing agents. **(A)** Western blot analysis of *PAN2* knockdown in HEK293A cells stably transfected with Dharmacon pGIPZ shRNA constructs (control or *PAN2*-specific) followed by puromycin selection and GFP-positive FACS sorting (see Methods). Lysates were immunoblotted with anti-*PAN2* antibody (Proteintech 16427-1-AP) and anti-actin (loading control). Pan2 (∼88 kDa) and a non-specific band are indicated. Quantification (right): Pan2/Actin ratio normalized to control; the Pan2KD population shows ∼75% reduction relative to control. Mean ± SD; ***p < 0.001 by two-tailed Student’s t-test. Data: 3 biological replicates. **(B)** MTT viability assay schematic. HEK293A cells were plated at 1,000 cells/well; drug or H₂O/DMSO vehicle was added 16 h post-plating; MTT reagent was added at 0/24/48/72 h post-treatment with absorbance read at 562 nm. **(C)** MTT viability response to nocodazole (0–80 nM). Left: absorbance at 562 nm vs. days post-treatment for control (blue) and Pan2KD (orange) cells across nocodazole concentrations. Right: normalized day-3 absorbance as percent of untreated, plotted as bars (mean ± SD). Pan2KD cells show significantly reduced viability relative to controls at 60 nM and 80 nM nocodazole; non-significant differences at 0, 20, and 40 nM. n.s., not significant; **p < 0.01, ****p < 0.0001 by multiple two-tailed Student’s t-tests (one per concentration). Data: 6 technical replicates per experiment, 3 biological replicates. **(D)** MTT viability response to colchicine (0–80 nM), layout as in (C). Pan2KD cells show significantly reduced viability relative to controls at all tested concentrations (10, 20, 40, and 80 nM). n.s., not significant; *p < 0.05, ****p < 0.0001 by multiple two-tailed Student’s t-tests (one per concentration). Data: 6 technical replicates per experiment, 3 biological replicates.

Both control and Pan2KD cells showed dose-dependent loss of viability upon treatment with nocodazole or colchicine over the time course (Figure 7C and 7D, left panels). At the 72-hour endpoint, Pan2KD cells were significantly more sensitive than controls to nocodazole at 60 and 80 nM (Figure 7C, right panel), and to colchicine at all tested concentrations (10, 20, 40, and 80 nM; Figure 7D, right panel). Thus, as in yeast, *PAN2* depletion sensitizes mammalian cells to microtubule destabilization.

### Pan2KD cells exhibit extended metaphase and multipolar spindles under colchicine treatment

To characterize the mitotic defects underlying the increased drug sensitivity of Pan2KD cells, we performed live-cell imaging of control and Pan2KD HEK293A cells labeled with CellLight™ Tubulin-RFP BacMam 2.0 under control and low-dose-colchicine (10 nM) conditions. Cells were seeded at 7,500 per well, drug was added 16 hours after plating, and imaging was performed continuously over 24 hours to capture spindle morphology and the durations of mitotic phases (Figure 8A). In the absence of colchicine, both control and Pan2KD cells progressed through the cell cycle with comparable metaphase duration and spindle morphology (Figure 8B; SFile 8).

**Figure 8:**
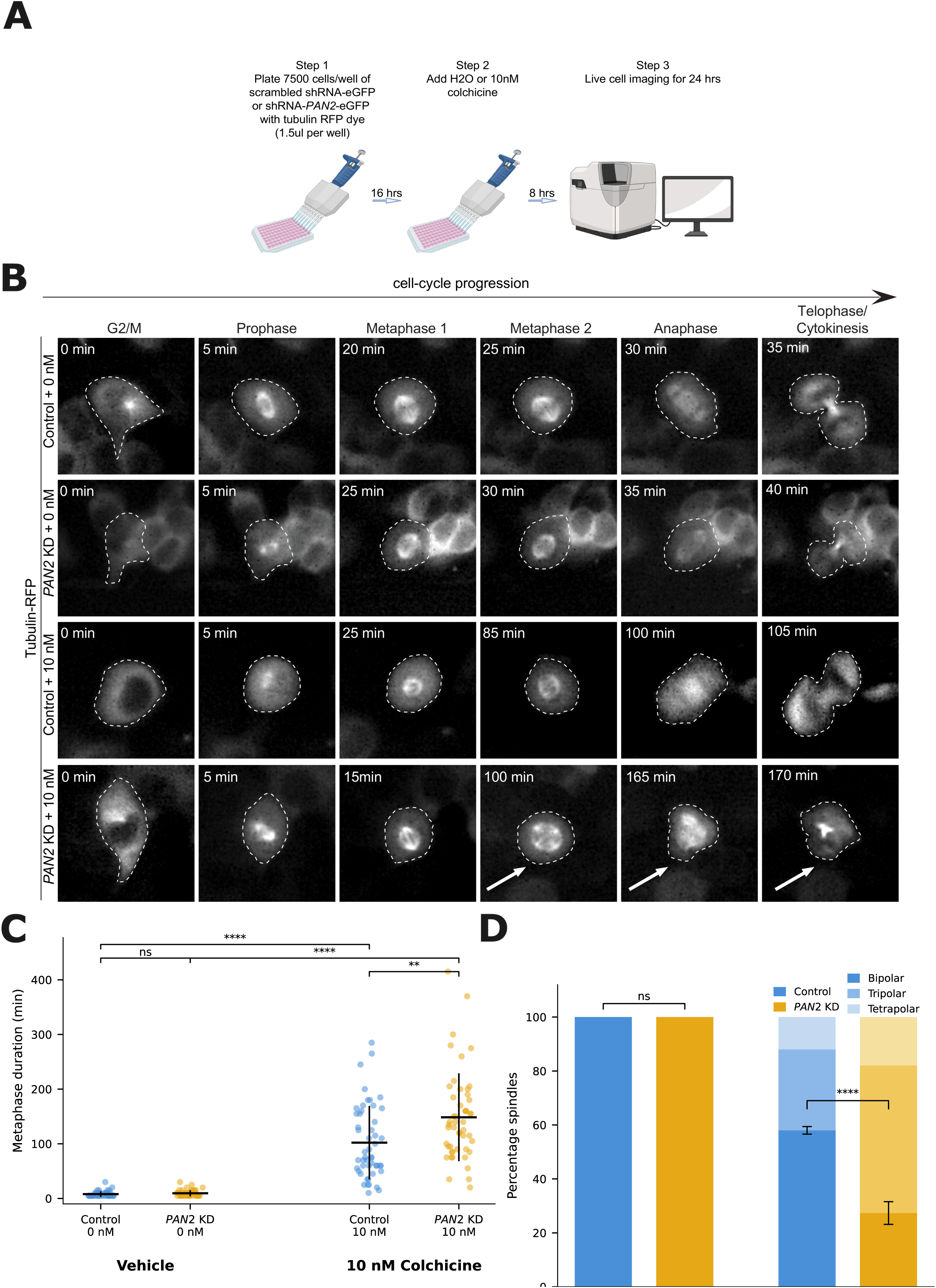
*PAN2* depletion impairs mitotic progression in human cells. **(A)** Experimental schematic. HEK293A cells stably expressing scrambled or *PAN2*-targeting Dharmacon pGIPZ shRNA-GFP constructs were plated at 7,500 cells/well, labeled with CellLight™ Tubulin-RFP BacMam 2.0 (1.5 μl/well), treated with H₂O vehicle or 10 nM colchicine after 16 h, and imaged live for 24 h beginning 8 h post-treatment. **(B)** Representative time-lapse image montage of control or Pan2KD HEK293A cells labeled with tubulin-RFP, cultured ± 10 nM colchicine. Images span cell-cycle progression from G2/M through cytokinesis (G2/M → prophase → metaphase 1 → metaphase 2 → anaphase → telophase/cytokinesis). In the absence of colchicine, both genotypes complete mitosis with comparable kinetics (∼35 min for control, ∼40 min for Pan2KD) and bipolar spindles. Upon 10 nM colchicine, control cells show prolonged but ultimately successful mitosis (∼105 min) with bipolar spindles, whereas Pan2KD cells display markedly prolonged metaphase (∼170 min) and multipolar spindles (arrows indicate aberrant pole organization). Time stamps (top left, in minutes). Scale bar = 10 μm. **(C)** Quantification of metaphase duration in control vs. Pan2KD cells, ± 10 nM colchicine. Each point = one cell; mean ± SD overlaid. Without colchicine, both genotypes complete metaphase rapidly. With 10 nM colchicine, both genotypes show prolonged metaphase, but Pan2KD cells exhibit significantly longer duration than control. n.s., not significant; **p < 0.01, ****p < 0.0001 by two-tailed Welch’s t-test. Data: 50 cells scored per condition from 3 biological replicates. **(D)** Spindle polarity classification at metaphase. Cells were scored as bipolar (2-pole), tripolar (3-pole), or tetrapolar (4-pole), and proportions are plotted as stacked bars per condition. Bar segments are colour-coded by genotype and pole-count: Control cells in shades of blue and Pan2KD cells in shades of yellow, with darker shades indicating bipolar and progressively lighter shades indicating tripolar and tetrapolar. Without colchicine, both genotypes form 100% bipolar spindles. With 10 nM colchicine, Control cells retain a majority of bipolar spindles (∼60% bipolar; ∼40% multipolar, defined as tripolar plus tetrapolar combined), whereas Pan2KD cells shift to a predominantly multipolar configuration (∼25% bipolar; ∼75% multipolar). n.s., not significant; ****p < 0.0001 by Fisher’s exact test (genotype × polarity-class) on raw cell counts. Data: 50 cells scored per condition from 2 biological replicates. See also SFiles 8 and 9.

Upon 10 nM colchicine treatment, control cells showed prolonged but ultimately successful mitosis, whereas Pan2KD cells spent markedly longer in metaphase (Figure 8B; SFile 9), quantified as a significant increase in metaphase duration relative to control (Figure 8C) and had a substantially higher propensity to form multipolar spindles (∼75% in Pan2KD vs. ∼40% in control; Figure 8D). Taken together, these findings indicate that Pan2 is required for spindle integrity during microtubule stress in mammalian cells, recapitulate the yeast phenotypes, and account for the reduced viability of Pan2KD cells under nocodazole and colchicine treatment.

## DISCUSSION

Our results define a stress-conditional role for PAN2-PAN3 deadenylase activity in mitotic robustness. In yeast, the requirement is uncovered only when microtubules are compromised: *pan2Δ* cells proliferate normally in unperturbed conditions but fail to complete mitosis under nocodazole, exhibiting G2/M arrest, disorganized spindles, and elevated cell death. The catalytic-dead Pan2-D1020A variant phenocopies *pan2Δ*, ruling out a scaffolding contribution and establishing that catalytic activity is the relevant function. In HEK293A cells, *PAN2* knockdown likewise produces extended metaphase, multipolar spindles, and reduced viability under nocodazole or colchicine, establishing that this function of PAN2-PAN3 is conserved from yeast to humans.

Why is PAN2-PAN3 catalytic activity dispensable under normal growth but essential when microtubules are compromised? Recent work has shown that bulk cytoplasmic deadenylation is suppressed during mitosis to stabilize the transcriptome of dividing cells, an effect attributed primarily to CCR4-NOT (Khalizeva *et al*., 2025). Our data argue that PAN2-PAN3 occupies a complementary role: rather than contributing to mitotic transcriptome stabilization, its catalytic activity is required when mitosis is challenged, suggesting a specialized stress-response module operating in parallel to the dominant deadenylation pathway. Stress-conditional genetic interactions of this kind are characteristic of buffering systems that maintain cellular robustness when other safeguards are exhausted (Hartman *et al*., 2001).

Consistent with cell-cycle-dependent regulation of PAN, multiple lines of evidence implicate CDK-mediated phosphorylation. In yeast, residue T415 in the Pan2 linker domain is a *CDK1* substrate (Holt *et al*., 2009). In mammalian cells, phosphorylation within the intrinsically disordered region adjacent to the PAM2 motif of Pan3 reduces its interaction with *PABPC1* and dampens PAN activity (Huang *et al*., 2013). Consistent with these biochemical observations, our SGA screen identified an alleviating genetic interaction between *PAN2* and *PHO85*, the yeast cyclin-dependent kinase that orchestrates stress and cell-cycle responses (SFile 2). The reduced fitness we observe upon Pan2 overexpression (Figure 1C, 6D) may likewise reflect a requirement to limit PAN activity outside its appropriate cell-cycle window.

If cell-cycle CDKs regulate PAN activity upstream, the downstream targets of PAN deadenylation under stress likely include specific cell-cycle mRNAs. RNA-seq comparing cells expressing wild-type versus catalytically inactive Pan2 under nocodazole identified a focused set of differentially expressed genes, including the mitotic cyclins *CLB1* and *CLN3*. However, neither cyclin is a clean phenotype driver: *CLB1* has the right function (G2/M cyclin) but the wrong direction (mRNA decreased rather than stabilized in catalytic-dead cells, opposite to a direct deadenylation target), while *CLN3* has the right direction (increased, consistent with deadenylation) but the wrong function (G1/S, disconnected from the spindle phenotype). Consistent with this, neither *clb1Δ* nor *cln3Δ* in a *pan2Δ* background modifies nocodazole sensitivity, though *PAN2* overexpression interacts aggravatingly with both deletions (Figure S6C), placing these cyclins within the PAN-dependent stress response as likely indirect effectors of cell-cycle dynamics.

Independent evidence supports an additional link to the mitotic apparatus: a whole-genome siRNA screen in HeLa cells reported *PAN2* loss as causing irregular nuclear shape and chromosome mis-segregation, with Pan2 and Pan3 found in a soluble G2 complex containing Bub1, Mad1, Ndc80, Nuf2, and microtubule motors (Cai *et al*., 2018; Neumann *et al*., 2010). Together with the genetic interactions between *pan2Δ* and tubulin-folding genes (*CIN1, CIN2, CIN4, GIM3, GIM4, PAC2*), and parallels to deadenylase-dependent control of tubulin homeostasis through the *TTC5*–CCR4-NOT axis (Batiuk *et al*., 2024; Rissland, 2024), these data place PAN2-PAN3 within a broader network coupling poly(A) tail control to spindle integrity.

We do not directly identify PAN2-PAN3 substrates in this work; resolving whether *CLB1*, *CLN3*, tubulin transcripts, or other mRNAs are deadenylation targets, and whether substrate specificity is conferred by recently described RNA-binding adaptors (Tang *et al*., 2025), will require poly(A)-tail-resolved sequencing approaches such as TAIL-seq or Nano3P-seq (Chang *et al*., 2014; Begik *et al*., 2023). The mammalian conservation of this requirement has clinical relevance. Microtubule-destabilizing agents and microtubule-stabilizing taxanes are mainstays of cancer chemotherapy, and tumor responses are often limited by cell-intrinsic resistance mechanisms. Our finding that PAN2-PAN3 is selectively required for survival under microtubule stress, but dispensable under basal proliferation, identifies it as a candidate vulnerability that could synergize with these agents while sparing unperturbed tissue. More broadly, the work positions PAN2-PAN3 deadenylation not as a constitutive layer of mRNA turnover but as a stress-responsive node of mitotic control, mirroring its emerging recognition as a tunable rheostat in development, immunity, and proliferative homeostasis.

## Supporting information

Yeast-codon-optimized human PAN2 sequence (FASTA).

Movie: WT yeast cell budding and dividing under DMSO (Tub1-yomWasabi).

Movie: WT yeast cell budding and dividing under nocodazole (Tub1-yomWasabi).

Movie: HEK293A control cell with bipolar spindle formation under colchicine.

Movie: HEK293A PAN2-knockdown cell with multipolar spindle formation under colchicine.

## ACKNOWLEDGEMENTS

We thank members of the Loewen and Roskelley labs for critical discussions. We thank Dr. Phil Hieter for sharing yeast overexpression plasmids and Dr. J. Thomas Beatty for sharing DH5α λpir *E. coli* cells. We thank the following core facilities at UBC: LSI imaging facility, ubcFLOW, and School of Biomedical Engineering (SBME) Sequencing Core (formerly BRC-Seq). Research was funded by the Canadian Institutes of Health Research (PJT-152967) and CFI Innovation Fund (2020) #39914 awarded to CJRL. This study was partially funded by the Canadian Institutes of Health Research (CIHR 169111) awarded to CDM and Canadian Institutes of Health Research (PJT-162253) awarded to LJH.

## AUTHOR CONTRIBUTIONS

J. V. conceived studies, collected samples, conducted experiments, interpreted experiments, wrote and edited the manuscript.

Z. H. collected and analyzed live-cell imaging data for mammalian cells.

J. A. R. B. collected and analyzed the flow-cytometry data for cell-cycle analysis in yeast cells.

P. D. assisted in designing the mammalian cell experiments and confirmed HEK293A *PAN2* KD GFP positive cells by flow cytometry.

B. P. Y. assisted in designing the yeast cell experiments and performed the initial testing of sensitivity of *pan2Δ* to nocodazole.

S. F. performed the bioinformatics analysis on RNA-seq data.

L. J. H. acquired the funding for cell-cycle analysis in yeast cells.

C. D. M. edited the manuscript and acquired funding for the microscope used for live-cell imaging data for mammalian cells.

C. D. R. conceived studies, interpreted experiments, edited the manuscript and acquired funding.

C. J. R. L. conceived studies, interpreted experiments, wrote and edited the manuscript, and acquired funding.

## DECLARATION OF GENERATIVE AI AND AI-ASSISTED TECHNOLOGIES IN THE WRITING PROCESS

During the preparation of this work the authors used Claude (Anthropic) in order to improve the readability and language of the manuscript. After using this tool, the authors reviewed and edited the content as needed and take full responsibility for the content of the publication.

## SUPPORTING FILE LEGENDS

**SFile 1:** Word file containing sequence of human *PAN2* codon optimized for yeast. Related to **Figure 1**.

**SFile 2:** Excel file containing Synthetic Genetic Analysis (SGA) data for *PAN2* in presence or absence of nocodazole. Related to **Figure 2**.

**SFile 3:** Excel file containing RNA-seq data for *pan2Δ* + WT Pan2 and *pan2Δ* + D1020A Pan2 in presence of nocodazole. Related to **Figure 6**.

**SFile 4:** Movie of a yeast wild-type (WT) cell budding and division upon treatment with DMSO. Tub1 is fluorescently tagged. Related to **Figure 4**.

**SFile 5:** Movie of a yeast wild-type (WT) budding and division upon treatment with nocodazole. Tub1 is fluorescently tagged. Related to **Figure 4**.

**SFile 6:** Movie of a yeast *pan2Δ* cell budding and division upon treatment with DMSO. Tub1 is fluorescently tagged. Related to **Figure 4**.

**SFile 7:** Movie of a yeast *pan2Δ* cell undergoing aberrant cell-cycle progression upon treatment with nocodazole, with budding but failed cell division. Tub1 is fluorescently tagged. Related to **Figure 4**.

**SFile 8:** Movie of a HEK293A control cell undergoing normal cell division (bipolar spindle formation) upon treatment with colchicine imaged for 24 h beginning 8 h post-treatment. Related to **Figure 8**.

**SFile 9:** Movie of a HEK293A *PAN2* KD cell undergoing abnormal cell division (multipolar spindle formation) upon treatment with colchicine imaged for 24 h beginning 8 h post-treatment. Related to **Figure 8**.

## STAR★METHODS

## KEY RESOURCES TABLE

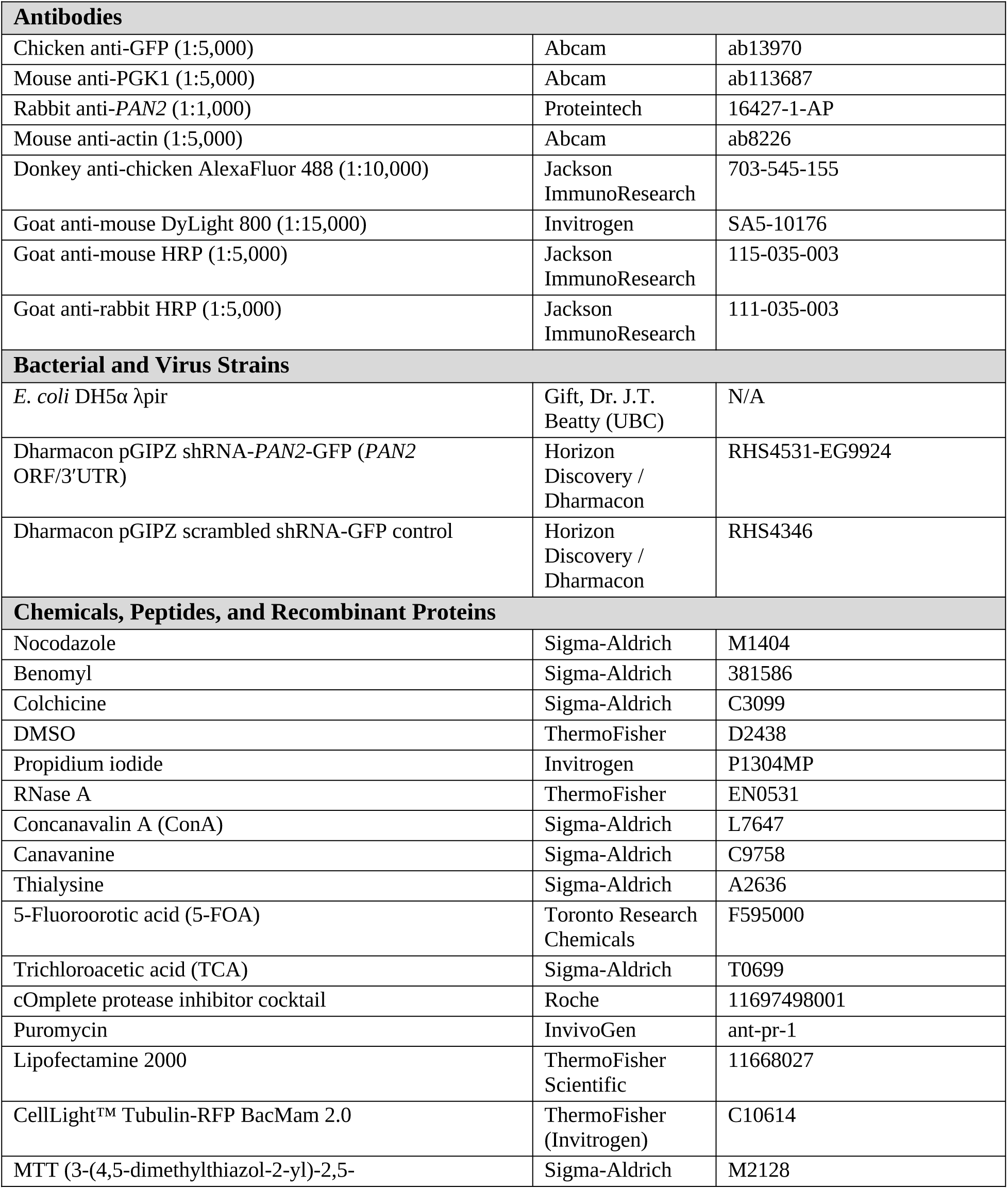
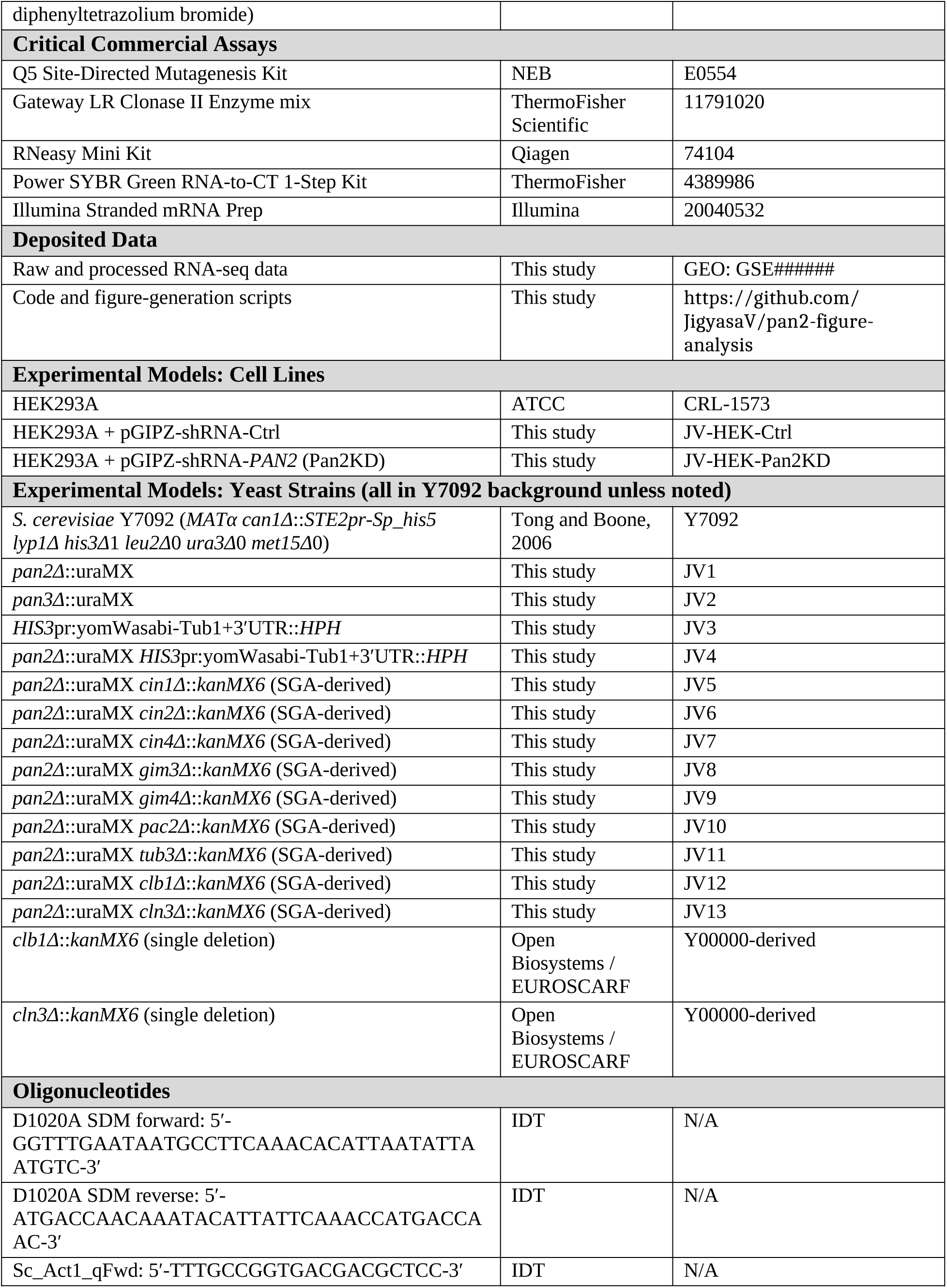
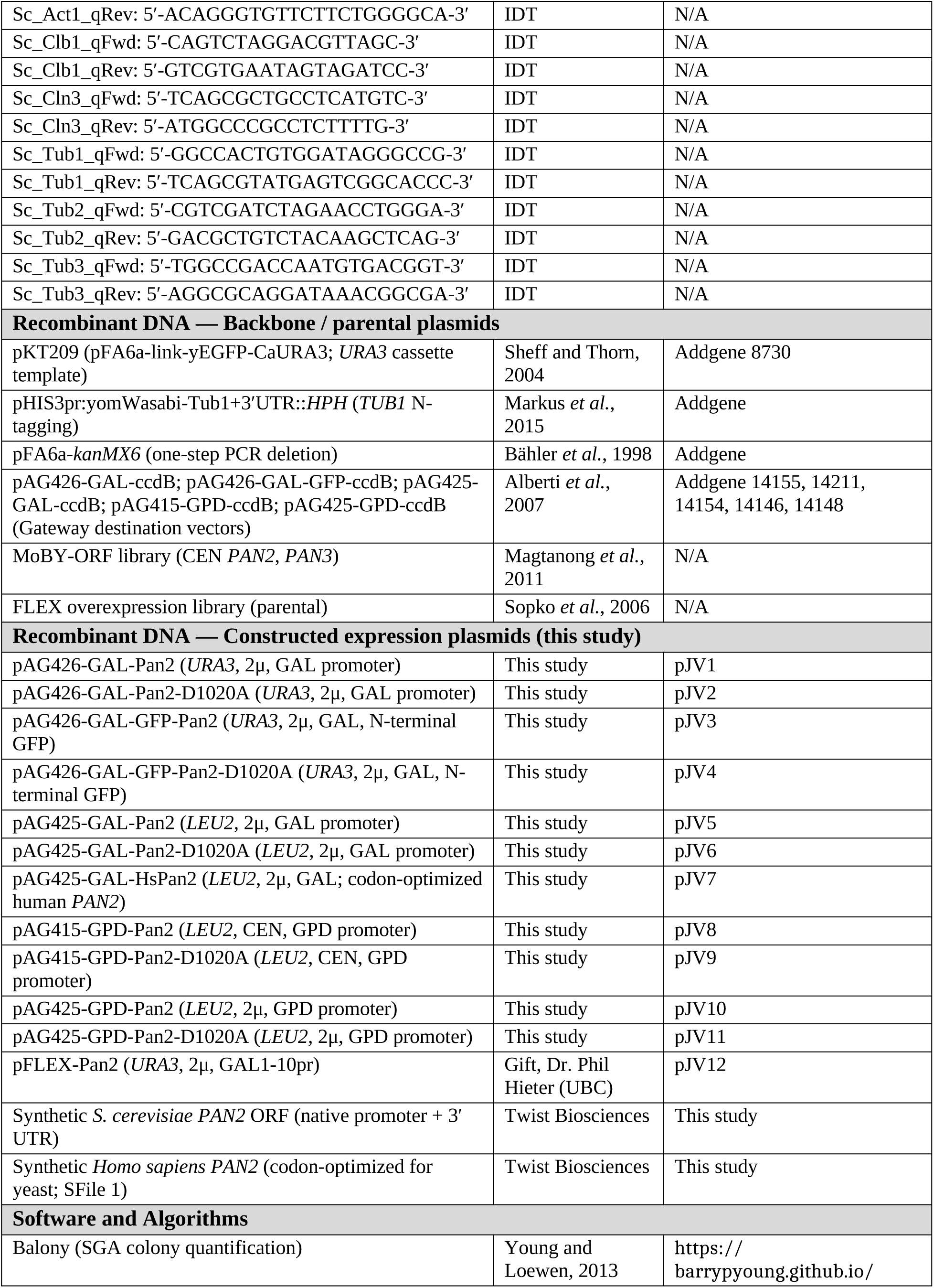
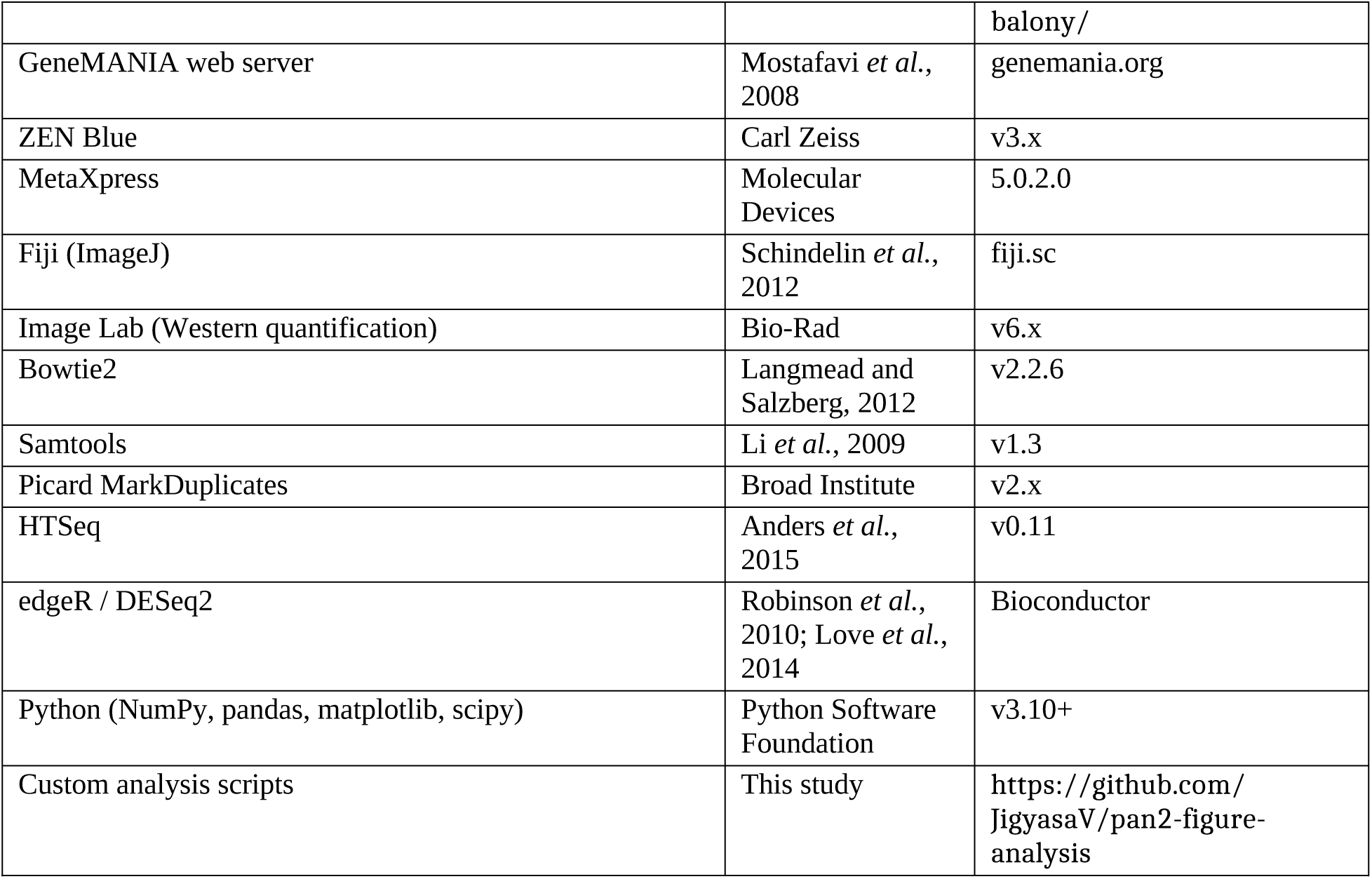

## RESOURCE AVAILABILITY

### Lead Contact

Further information and requests for resources and reagents should be directed to and will be fulfilled by the Lead Contact, Christopher J.R. Loewen.

### Materials Availability

All yeast strains and plasmids generated in this study are available from the Lead Contact upon request without restriction. Mammalian cell lines and shRNA constructs are available upon request, subject to the corresponding material transfer agreements with Horizon Discovery / Dharmacon.

Raw RNA-sequencing reads and processed count matrices have been deposited at NCBI Gene Expression Omnibus (GEO) under accession GSE###### (to be released upon publication).

Original imaging data, qRT-PCR raw Ct values, MTT raw absorbance, and SGA colony-size data are available from the Lead Contact upon request. All code used for figure generation and statistical analysis is publicly available at https://github.com/JigyasaV/pan2-figure-analysis. Any additional information required to reanalyze the data reported in this paper is available from the Lead Contact upon request.

## EXPERIMENTAL MODEL AND SUBJECT DETAILS

### Yeast Strains

*Saccharomyces cerevisiae* strains used in this study were derived from BY4741 (*MATa his3Δ*1 *leu2Δ*0 *met15Δ*0 *ura3Δ*0), JHY716 and Y7092 (*MATα can1Δ*::*STE2pr*-*Sp_his5 lyp1Δ his3Δ*1 *leu2Δ*0 *ura3Δ*0 *met15Δ*0). Single and double deletion strains were constructed by homologous recombination of PCR-generated linear cassettes amplified from pKT209 (*URA3*), pFA6a-*kanMX6* (*KanMX*), or pHIS3p:yomWasabi-*TUB1*+3′UTR::*HPH* (*HPH*), with successful integration verified by colony PCR and, where applicable, Sanger sequencing. All strains are listed in the Key Resources Table.

Yeast strains were maintained on YPD agar (1% yeast extract, 2% peptone, 2% dextrose) for general propagation, and on synthetic complete (SC) media (Yeast Nitrogen Base, ammonium sulfate, complete amino-acid supplement, 2% dextrose) with appropriate amino-acid dropouts for plasmid selection. All yeast cultures were grown at 30 °C unless otherwise specified.

HEK293A cells (ATCC CRL-1573; female human embryonic kidney origin) were cultured in DMEM/F12 medium (Sigma) supplemented with 10% fetal bovine serum (FBS) and 1% penicillin–streptomycin in a humidified incubator at 37 °C and 5% CO2. Cells were passaged every 2–3 days at ∼80% confluency (ThermoFisher). Cell-line identity was confirmed by STR profiling and routine mycoplasma testing was performed. All experiments were performed within 15 passages of thawing.

## METHOD DETAILS

### Plasmids and Constructs

*Saccharomyces cerevisiae PAN2* and *PAN3* CEN-based complementation plasmids were obtained from the MoBY-ORF collection (Magtanong *et al*., 2011). Galactose-inducible overexpression plasmids were obtained from the FLEX overexpression library (Sopko *et al*., 2006), generously provided by Tejomayee Singh from the laboratory of Dr. Phil Hieter (UBC). The complete *S. cerevisiae PAN2* coding sequence with native promoter and 3′ UTR, as well as a yeast-codon-optimized *Homo sapiens PAN2* sequence (provided in SFile 1), were synthesized by Twist Biosciences and cloned into Gateway destination vectors (pAG426GAL-ccdB, pAG426-GAL-GFP-ccdB, pAG425-GAL-ccdB, pAG415-GPD-ccdB, pAG425-GPD-ccdB; Alberti *et al*., 2007) using the Gateway LR Clonase II kit (ThermoFisher). For GFP-tagged constructs, GFP was fused to the N-terminus of Pan2 (yielding GFP-Pan2). Plasmid identity was confirmed by Nanopore sequencing (Plasmidsaurus).

The deadenylase-dead Pan2-D1020A point mutation was introduced into the *PAN2* coding sequence by site-directed mutagenesis using the Q5 Site-Directed Mutagenesis Kit (NEB E0554) with the primer pair 5′-GGTTTGAATAATGCCTTCAAACACATTAATATTAATGTC-3′ (forward) and 5′-ATGACCAACAAATACATTATTCAAACCATGACCAAC-3′ (reverse). PCR products were transformed and recombined in *E. coli* DH5α λpir (gift of Dr. J.T. Beatty, UBC), and verified by Sanger sequencing.

For C-terminal tubulin tagging in JHY716, the N-terminal yomWasabi-Tub1 fusion (with native 3′UTR) cassette strategy was used as described in Markus *et al*., 2015. Briefly, the parental yomWasabi-Tub1 tagging plasmid was modified to include an additional 618 nucleotides corresponding to the genomic region downstream of the *TUB1* locus, extending past both potential polyadenylation signals (AATAAA at +145 nt; ATTAAA at +540 nt downstream of the stop codon). The modified plasmid pHIS3p:yomWasabi-*TUB1*+3′UTR was linearized with BsaBI prior to transformation. Following integration, both the endogenous untagged *TUB1* and the integrated tagged copy retain the native 3′UTR sequence.

### Yeast Liquid Growth Assays

Yeast strains were grown overnight in liquid SC media supplemented with appropriate amino acids, diluted into fresh medium, and grown to logarithmic phase (OD600 0.2–0.5). Cultures were normalized and diluted to a starting OD600 of 0.005–0.010 in fresh SC media containing the indicated concentrations of nocodazole (Sigma) or DMSO vehicle (≤0.5% v/v). 200 μL aliquots were dispensed into clear-bottom 96-well plates in technical hexaplicate (6 wells per condition). Plates were incubated at 30 °C with continuous orbital shaking at 600 rpm in a BioTek Epoch 2 Microplate Spectrophotometer with OD600 measured every 5 minutes for 24 hours. For each independent biological replicate, 6 technical replicates were averaged; three independent biological replicates were performed per experiment (total n = 6 technical × 3 biological).

For exponential growth-rate determination, OD600 values within the log-linear growth window (typically OD 0.3–1.5) were log-transformed, and growth constants (k, h⁻¹) were derived from the slope of ln(OD600) versus time using linear regression. Growth-rate assays under nocodazole treatment (in which growth was non-exponential) were instead reported as raw growth curves.

Statistical analyses were performed in Python (NumPy, pandas, and SciPy stats module) using custom scripts available at github.com/JigyasaV/pan2-figure-analysis. Two-group comparisons used Student’s t-test (scipy.stats.ttest_ind) or Welch’s t-test (equal_var=False) where indicated; multiple-group comparisons used one-way ANOVA with appropriate post-hoc corrections.

### Synthetic Genetic Array (SGA) Analysis

SGA screens were performed essentially as described (Tong and Boone, 2006; Young and Loewen, 2013). The SGA query strain was generated by integrating a *URA3* cassette amplified from pKT209 at the *PAN2* locus in Y7092. The *pan2Δ*::*URA3* query was crossed against the genome-wide non-essential haploid deletion collection (∼4,800 strains, marked with *kanMX6*) using a Singer RoToR HDA pinning robot. Diploids were selected and induced to sporulate.

*MATa* haploid double mutants were obtained by sequential selection on SC-His/Arg/Lys + 100 μg/mL canavanine + 100 μg/mL thialysine. Counter-selection on 5-FOA media was used to obtain control plates lacking the *pan2Δ* mutation; experimental plates were obtained by selection on SC-Ura. Both control and experimental plates were re-pinned onto identical media containing 1.5 μM nocodazole for the drug-treated screens.

Plates were incubated at 30 °C, photographed using a Canon CanoScan flatbed scanner, and colony sizes were quantified using Balony software (Young and Loewen, 2013). Each screen was performed in 3/3 biological replicates. Aggravating genetic interactions were called at a colony-size ratio cutoff of ≤ 0.7 (three standard deviations from the diagonal; 99.7% CI) at p < 0.05 across all replicates; alleviating interactions used a cutoff of ≥ 1.3 with the same significance criteria. Gene Ontology enrichment and protein–protein interaction network analysis was performed using the GeneMANIA web server (Mostafavi *et al*., 2008) with Benjamini–Hochberg FDR-adjusted p-values (q-values) from hypergeometric tests.

### Western Blotting

Yeast lysates were prepared by trichloroacetic acid (TCA) precipitation. Briefly, 5 mL of culture at OD600 ≈ 0.8 was harvested by centrifugation, resuspended in 20% TCA, and lysed by vortexing with acid-washed glass beads for 3 minutes. Precipitated proteins were pelleted, washed, resuspended in 2× SDS-PAGE sample buffer, neutralized with Tris-HCl, and boiled.

Lysates were separated on 4–12% gradient SDS-PAGE gels and transferred to nitrocellulose (0.2 μm) by wet transfer. Membranes were blocked in 5% non-fat milk in TBST for 1 hour at room temperature and probed with chicken anti-GFP (1:5,000; Abcam ab13970) and mouse anti-PGK1 (1:5,000; Abcam ab113687, loading control) overnight at 4 °C. Detection used donkey anti-chicken AlexaFluor 488 (1:10,000; Jackson 703-545-155) and goat anti-mouse DyLight 800 (1:15,000; Invitrogen SA5-10176) and visualized on a Bio-Rad ChemiDoc MP system. Band intensities were quantified using Image Lab software (Bio-Rad).

Mammalian lysates were prepared by lysing HEK293A cells in RIPA buffer (50 mM Tris-HCl pH 7.4, 150 mM NaCl, 1% Triton X-100, 0.5% sodium deoxycholate, 0.1% SDS, 5 mM EDTA) supplemented with cOmplete protease inhibitor cocktail (Roche). Lysates were clarified by centrifugation at 14,000 × g for 10 minutes and quantified by Bradford assay. 20 μg of total protein per lane was separated by SDS-PAGE and transferred to nitrocellulose. Membranes were probed with rabbit anti-*PAN2* (1:1,000; Proteintech 16427-1-AP) and mouse anti-actin (1:5,000; Abcam ab8226), followed by HRP-conjugated goat anti-rabbit (1:5,000; Jackson 111-035-003) and goat anti-mouse (1:5,000; Jackson 115-035-003) secondaries. Detection was by ECL chemiluminescence; band intensities were quantified by densitometry using Image Lab.

### FACS Analysis of DNA Content

Yeast cells cultured in SC media ± 6 μM nocodazole for 3, 6, and 10 hours were harvested by centrifugation, washed with sterile water, and fixed in ice-cold 70% ethanol overnight at 4 °C. Fixed cells were rehydrated in PBS, treated with RNase A (100 μg/mL; ThermoFisher) for 3 hours at 37 °C, and stained with propidium iodide (5 μg/mL; Invitrogen) for 1 hour at room temperature. Cells were analyzed on a Beckman Coulter CytoFLEX LX cytometer with 488-nm excitation and 610/20-nm bandpass detection. Doublets and debris were excluded by forward-scatter (FSC) versus side-scatter (SSC) gating, and ≥10,000 single-cell events were collected per sample. Three independent biological replicates were analyzed per condition. Mean FSC intensity was used as a proxy for cell volume. Data were analyzed in FlowJo v10.

### Live-Cell Imaging of Yeast

Live-cell imaging was performed in 96-well glass-bottomed microscopy plates (Cellvis P96-1.5H-N) coated with Concanavalin A (ConA; Sigma L7647). A 10 mg/mL ConA stock was prepared, aliquoted, and stored at −20 °C; a 1 mg/mL working solution was prepared in water and stored at 4 °C. ConA solution was applied to each well, incubated for 5 minutes at room temperature, aspirated, and dried for 5 minutes.

JHY716-derived *TUB1*-yomWasabi yeast strains were grown overnight in sterile-filtered SC media and diluted to log phase the next morning. 20 μL of cell suspension was applied to each ConA-coated well; after 10 minutes of incubation to allow adhesion, the medium was removed and replaced with 40 μL of fresh SC media (without cells) containing 6 or 12 μM nocodazole or DMSO vehicle.

Imaging was performed on a Zeiss CellDiscoverer 7 system equilibrated to 30 °C, using a Plan-Apochromat 50×/1.2 water-immersion objective and a 1× tubelens for a final magnification of 50× (effective NA 1.2). Images were acquired in brightfield (TL LED lamp) and GFP (LED module 470 nm) channels at 5-minute intervals for 480 minutes (8 hours) using definite focus. Z-stacks of 9 planes spaced ∼0.125 μm apart were acquired at each timepoint. Two fields of view were imaged per well at non-overlapping locations; 2 biological replicates were imaged per condition.

Image processing was performed in ZEN Blue (Carl Zeiss). Z-stacks were deconvolved using a constrained iterative algorithm and projected using extended depth of focus to an effective depth of ∼1 μm. Cell segmentation was performed on the GFP channel using Gaussian smoothing (σ = 2) followed by iso-data thresholding for light regions. For each segmented cell, area and mean GFP intensity were measured. Cells with mean GFP intensity > 8000 (an empirical threshold corresponding to lysed cells with elevated cytoplasmic GFP signal) were classified as dead. For each well, total cell counts, dead-cell counts, and cell areas were compiled across the imaging time course.

### Endpoint Fluorescence Imaging and Microtubule Organization Quantification

For quantitative endpoint analysis of microtubule organization, wild-type and *pan2Δ* cells expressing *TUB1*-yomWasabi were grown in shaking liquid SC media supplemented with 6 μM nocodazole or DMSO vehicle for 10 hours at 30 °C, then loaded onto poly-lysine-coated 8-well glass-bottomed slides (Lab-Tek II) for imaging. Imaging was performed on the same Zeiss CellDiscoverer 7 system used for live-cell imaging (described above), with a Plan-Apochromat 50×/1.2 water-immersion objective (1× tubelens; final magnification 50×, NA 1.2), at 30 °C. Brightfield (TL LED lamp) and GFP (LED module 470 nm) channels were acquired. Single-timepoint Z-stacks (11 planes, 0.5 μm spacing) were acquired and projected as maximum-intensity projections. Dead cells were excluded from analysis based on loss of GFP fluorescence and morphological criteria (loss of turgor).

For each segmented cell (region of interest, ROI), peak-to-mean intensity ratio was calculated as the maximum pixel intensity divided by the mean pixel intensity within the ROI, capturing the prominence of bright tubulin puncta relative to diffuse cytoplasmic signal. The coefficient of variation (CV) of GFP intensity was calculated as the standard deviation of pixel intensities within the ROI divided by the mean intensity, providing a measure of fluorescence-distribution heterogeneity. Cell-boundary segmentation and per-cell area, mean intensity, and intensity-distribution metrics were computed using a custom Python pipeline (scikit-image watershed segmentation; SciPy ndimage; pandas) (van der Walt *et al*., 2014) available at github.com/JigyasaV/pan2-figure-analysis. For kernel density estimation, distributions of peak-to-mean ratio, CV, and cell area were computed using the Gaussian kernel in Python (scipy.stats.gaussian_kde) with automatic bandwidth selection. ≥4,000 individual cells per condition were pooled across 3 independent biological replicates. Statistical comparisons were performed by two-tailed Welch’s t-test with Bonferroni correction (three planned contrasts; *p < 0.05, **p < 0.01, ***p < 0.001, ****p < 0.0001).

### RNA Isolation and qRT-PCR

Yeast cells cultured ± 6 μM nocodazole for 6 hours in SC media were harvested by centrifugation. Total RNA was isolated using the RNeasy Mini Kit (Qiagen 74104) with on-column DNase I digestion according to the manufacturer’s protocol. RNA concentration and purity were assessed by spectrophotometry (NanoDrop); integrity was verified by agarose gel electrophoresis. Quantitative reverse-transcriptase PCR was performed using the Power SYBR Green RNA-to-CT 1-Step Kit (ThermoFisher 4389986) on an Applied Biosystems StepOnePlus Real-Time PCR System. Cycling conditions were: 48 °C 30 min (RT step); 95 °C 10 min; 40 cycles of 95 °C 15 s and 60 °C 60 s; melt curve from 60–95 °C at 0.1 °C/s. Primer pairs were validated to have efficiencies of 95–105% by standard curves. Relative expression was calculated by the ΔΔCt method with *ACT1* as the endogenous control and *pan2Δ* + Pan2-WT + DMSO as the calibrator. Fold-change values were calculated as 2^(−ΔΔCt). Three independent biological replicates were analyzed, each with technical triplicates. Statistical analysis used pairwise two-tailed Student’s t-tests (scipy.stats.ttest_ind) implemented in Python; *p < 0.05, **p < 0.01, ***p < 0.001, ****p < 0.0001. Analysis scripts are available at github.com/JigyasaV/pan2-figure-analysis.

### RNA Sequencing and Bioinformatics

Duplicate cultures of *pan2Δ* + Pan2-WT and *pan2Δ* + Pan2-D1020A were grown in SC media at 30 °C to OD600 ≈ 0.6, then treated with 6 μM nocodazole for 6 hours. Cells were harvested by centrifugation, snap-frozen in liquid nitrogen, and total RNA was extracted using the RNeasy Mini Kit (Qiagen) with on-column DNase I digestion. RNA quality was assessed by Agilent 2100 Bioanalyzer; only samples with RIN > 8 were used.

Sequencing libraries were prepared using the Illumina Stranded mRNA Prep kit with poly(A) enrichment according to the manufacturer’s protocol. Library quality and quantity were verified by Bioanalyzer and Qubit fluorometry, respectively. Libraries were pooled in equimolar amounts and sequenced on an Illumina NextSeq2000 in paired-end 59 × 59 bp configuration at the BRC Sequencing Facility (UBC), generating ∼30 million read pairs per sample.

Reads were aligned to the *S. cerevisiae* reference genome and transcriptome (R64-1-1, Ensembl Release 107) using Bowtie2 (v2.2.6). Read counts per annotated gene were generated using HTSeq (v0.11) in strand-specific mode against the Ensembl GTF. Multiple differential-expression pipelines (edgeR and DESeq2) were applied independently and results were intersected. Differentially expressed genes were called using a z-score threshold of |z| > 2 (computed from the per-gene log2-fold-change distribution; equivalent to log2FC > +1.180 or < −1.055), p < 0.01, and log2 CPM > 1. Of 6,031 ORFs analyzed, 129 genes were upregulated and 39 downregulated in Pan2-D1020A relative to Pan2-WT under nocodazole. Gene Ontology enrichment analysis was performed using clusterProfiler (v4.0) testing for overrepresentation against the genome background; significance was assessed by hypergeometric test with Benjamini–Hochberg adjustment. Volcano plots were generated in Python (matplotlib).

### Generation of Stable *PAN2* Knockdown HEK293A Cell Populations

HEK293A cells were maintained in DMEM/F12 (Sigma) + 10% FBS at 37 °C, 5% CO2. To generate *PAN2* knockdown populations, cells were transfected with a Dharmacon pGIPZ shRNA set (RHS4531-EG9924) in which shRNAs targeting the *PAN2* ORF/3′UTR are co-expressed with GFP. Transfections used Lipofectamine 2000 (ThermoFisher 11668027) according to the manufacturer’s instructions; a non-targeting scrambled shRNA-GFP construct served as control.

Forty-eight hours post-transfection, puromycin selection was initiated at 1 μg/mL and maintained for 14 days to enrich for cells stably integrating the shRNA construct. Puromycin-resistant cells were sorted by fluorescence-activated cell sorting (FACS) on the basis of GFP expression to isolate stably transfected populations. *PAN2* knockdown efficiency was confirmed by Western blotting using rabbit anti-*PAN2* (Proteintech 16427-1-AP, 1:1,000), with knockdown estimated at ∼75% by densitometry relative to scrambled-shRNA control.

### MTT Viability Assays

HEK293A control and Pan2KD cells were seeded into clear-bottom 96-well plates at 1,000 cells per well in 100 μL DMEM/F12 + 10% FBS. After 16 hours, drug or vehicle was added to a final concentration of 0–80 nM nocodazole or 0–80 nM colchicine (in 0.5% DMSO vehicle). At 0, 24, 48, and 72 hours post-treatment, MTT (Sigma M2128) was added to wells at 1 mg/mL final concentration in growth media and incubated for 2 hours at 37 °C to allow formazan crystal formation. Medium was aspirated and crystals dissolved in 100 μL DMSO per well. Absorbance was read at 562 nm on a Tecan Sunrise microplate reader. For each condition, 6 technical replicates were performed; 3 independent biological replicates were completed.

Cell viability was calculated as (A562 treated / A562 untreated) × 100% per condition. Statistical comparisons between control and Pan2KD cells at each drug concentration were performed by multiple two-tailed Student’s t-tests (one per concentration) implemented in Python (scipy.stats.ttest_ind); *p < 0.05, **p < 0.01, ****p < 0.0001. Analysis scripts are available at github.com/JigyasaV/pan2-figure-analysis.

### Live-Cell Imaging of HEK293A Cells

For live-cell imaging of mitotic progression, HEK293A control and Pan2KD cells were seeded into 96-well clear-bottom plates (Corning) at 7,500 cells per well in DMEM/F12 + 10% FBS. CellLight™ Tubulin-RFP BacMam 2.0 reagent (ThermoFisher C10614) was added at 1.5 μL/well 24 hours prior to imaging to label microtubules. Cells were treated with 10 nM colchicine or H2O vehicle 16 hours after plating, and imaging began 8 hours post-treatment.

Imaging was performed on an ImageXpress Micro XL epifluorescence microscope (Molecular Devices) with a 37 °C, 5% CO2 environmental chamber, using a 40×/0.75 NA dry objective. Time-lapse images were acquired every 15 minutes over a continuous 24-hour window using 100-ms exposures at 2×2 binning and 25% lamp intensity per channel. Image sequences and movies were assembled in MetaXpress 5.0.2.0.

Metaphase duration was determined by manual frame-counting in the tubulin-RFP channel: metaphase was defined as the interval from chromosome alignment at the metaphase plate until anaphase onset (chromatid separation). 50 mitotic cells per condition were scored across 3 biological replicates. Spindle polarity was classified at metaphase as bipolar (2 poles), tripolar (3 poles), or tetrapolar (4 poles) by visual inspection; 50 cells per condition were classified across 2 biological replicates.

## QUANTIFICATION AND STATISTICAL ANALYSIS

All statistical analyses were performed in Python using NumPy, pandas, and the SciPy stats module. Custom analysis scripts (one per figure panel) are publicly available at github.com/JigyasaV/pan2-figure-analysis; in all cases p < 0.05 was considered statistically significant. Asterisks denote: *p < 0.05, **p < 0.01, ***p < 0.001, ****p < 0.0001; n.s., not significant.

| Statistical tests by figure |  |  |
| --- | --- | --- |
| Fig 1A/B (growth curves) | No formal test on curves; descriptive | n = 6 technical × 3 biological |
| Fig 1C/D (DMSO growth-rate bars) | Two-tailed Student's t-test | n = 3–6 wells per genotype |
| Fig 2B (double-mutant growth) | No formal test on curves; descriptive | n = 6 technical × 3 biological |
| Fig 3A (FACS DNA content) | Descriptive (representative histograms) | n = 3 biological, $\geq 10,000$ cells/sample |
| Fig 3B/C (cell diameter) | Two-tailed Welch's t-test | n > 100 cells/condition, 3 biological |
| Fig 4A/B (cell counts, dead cells) | Two-tailed Welch's t-test on endpoint values | n = 4 biological replicates |
| Fig 4C (cell area over time) | Two-tailed Welch's t-test on endpoint values | n = 380–2,726 cells/timepoint, 3 biological |
| Fig 5B–D (peak/mean, CV, area) | Two-tailed Welch's t-test, Bonferroni-corrected (3 planned contrasts) | n > 4,000 cells/condition, 3 biological |
| Fig 6A (RNA-seq differential expression) | edgeR + DESeq2 intersection; $\vert z\vert > 2$, $p < 0.01$, $\log_2\text{CPM} > 1$ | n = 2 biological replicates |
| Fig 6B (GO enrichment) | Hypergeometric test with Benjamini–Hochberg adjustment | n = 168 differentially expressed genes |
| Fig 6C (qRT-PCR fold-change) | Pairwise two-tailed Student's t-test | n = 3 biological × 3 technical |
| Fig 6D (Pan2 OE growth-rate bars) | Two-tailed Student's t-test | n = 6 technical × 3 biological |
| Fig 7A (Western quantification) | Two-tailed Student's t-test | n = 3 biological |
| Fig 7C/D (MTT dose-response) | Multiple two-tailed Student's t-tests (one per concentration) | n = 6 technical × 3 biological |
| Fig 8C (metaphase duration) | Two-tailed Welch's t-test | n = 50 cells/condition, 3 biological |
| Fig 8D (spindle polarity) | Fisher's exact test (genotype × polarity-class) | n = 50 cells/condition, 2 biological |
| SFig 2C (tubulin mRNA $\Delta\text{Ct}$) | Pairwise two-tailed Welch's t-test (n.s. across all 6 pairs/gene) | n = 3 biological × 3 technical |
| SFig 6A (RNA-seq replicate correlation) | Spearman correlation | n = 2 biological/genotype |
| SFig 6C (epistasis analysis) | Two-tailed Student's t-test (observed vs. multiplicative null) | n = 6 technical × 3 biological |

For all bar plots, error bars represent ± standard deviation (SD) unless otherwise noted; growth-curve shaded bands represent ± SD pooled across biological replicates, except Fig 4A/B which use ± 95% confidence interval (n = 4 biological replicates, t-distribution df = 3) and Fig 4C / SFig 4C / SFig 6B which use ± SEM. Sample sizes (“n”) denote the number of cells, technical replicates, and biological replicates as indicated in each figure legend.

## SUPPLEMENTARY FIGURE LEGENDS

**Figure S1:**
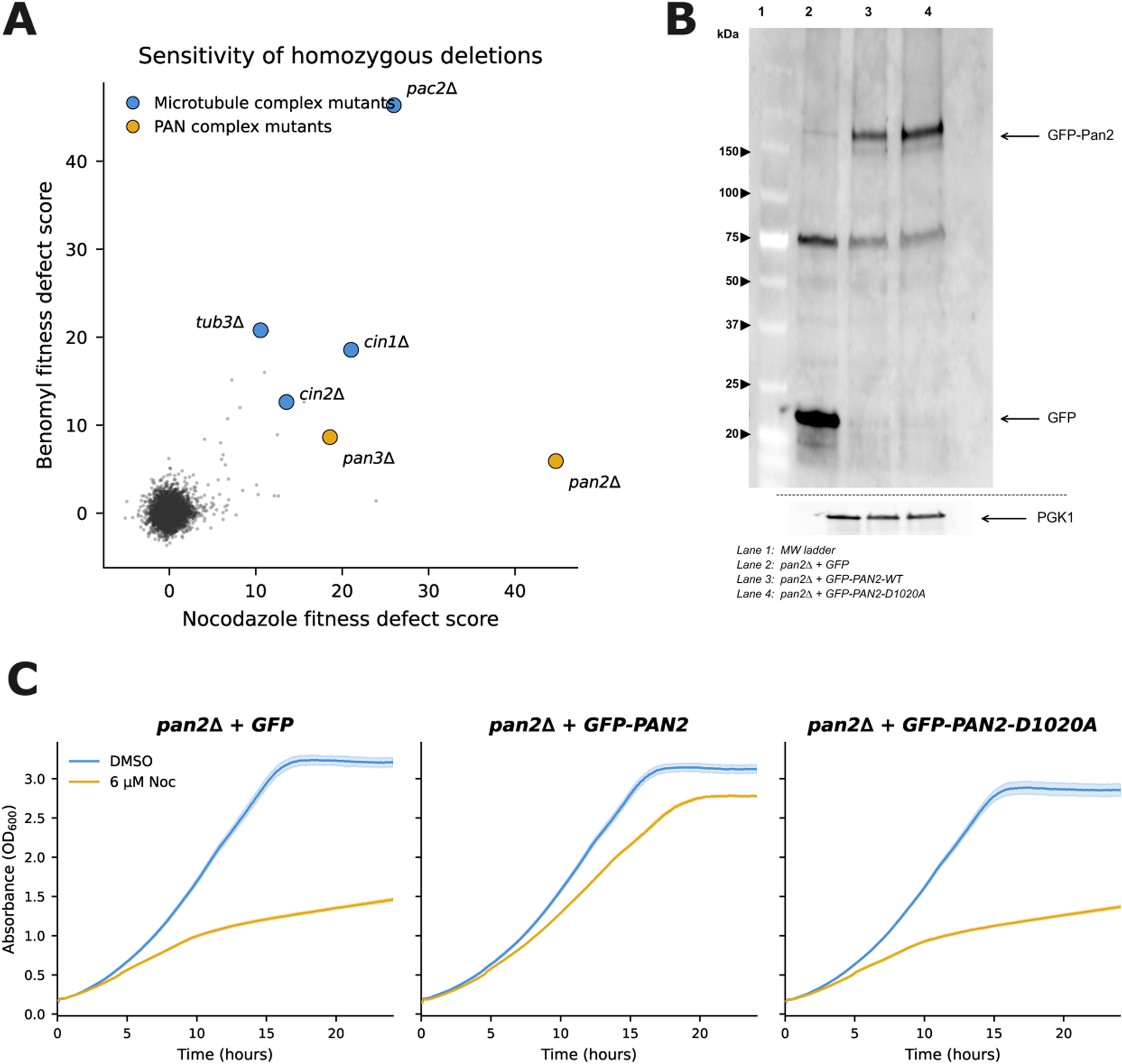
Sensitivity screen, GFP-Pan2 expression, and GFP-tagged complementation. **(A)** Sensitivity ranking of homozygous diploid yeast deletion strains to nocodazole (x-axis) and benomyl (y-axis), based on fitness defect scores from a genome-wide chemical-genetic screen of ∼4,800 mutants (Hillenmeyer *et al*., 2008; Lee *et al*., 2014). Both *pan2Δ* and *pan3Δ* (orange, PAN complex) rank in the top 20 most-sensitive strains for both drugs, alongside microtubule-pathway mutants *pac2Δ*, *cin1Δ*, *cin2Δ*, and *tub3Δ* (blue, microtubule complex). **(B)** Western blot validation of GFP-Pan2 expression. (1) MW ladder, lysates from *pan2Δ* cells expressing (2) GFP alone, (3) GFP-*PAN2*-WT, or (4) GFP-*PAN2*-D1020A from a centromeric plasmid, immunoblotted with anti-GFP antibody. Upper band: GFP-Pan2 (∼150 kDa); lower band: free GFP (∼25 kDa). PGK1 serves as loading control. Both fusion proteins are expressed at comparable levels, confirming that the catalytic-dead phenotype reflects loss of enzymatic activity rather than reduced protein abundance. **(C)** Complementation growth curves with GFP-tagged Pan2 variants. *pan2Δ* cells expressing GFP alone, GFP-*PAN2* (WT), or GFP-*PAN2*-D1020A cultured with 6 μM nocodazole (orange) or DMSO (blue) over 24 h at 30 °C. GFP-Pan2(WT) largely restores nocodazole resistance, whereas GFP-Pan2(D1020A) does not, confirming that rescue depends on Pan2 catalytic activity. Absorbance (OD₆₀₀) plotted with shaded ± SD. Data: 6 technical replicates per experiment, 3 biological replicates.

**Figure S2:**
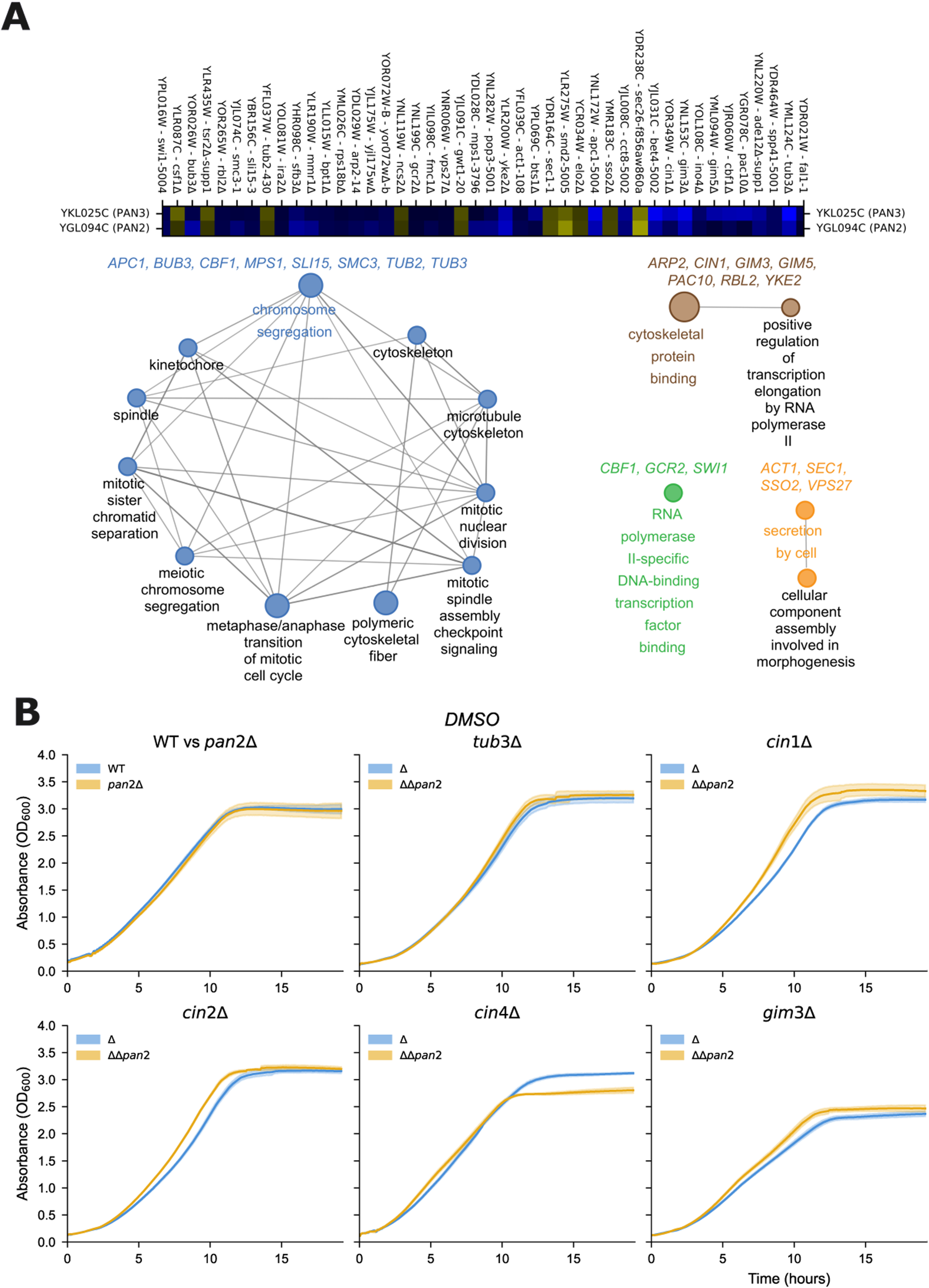
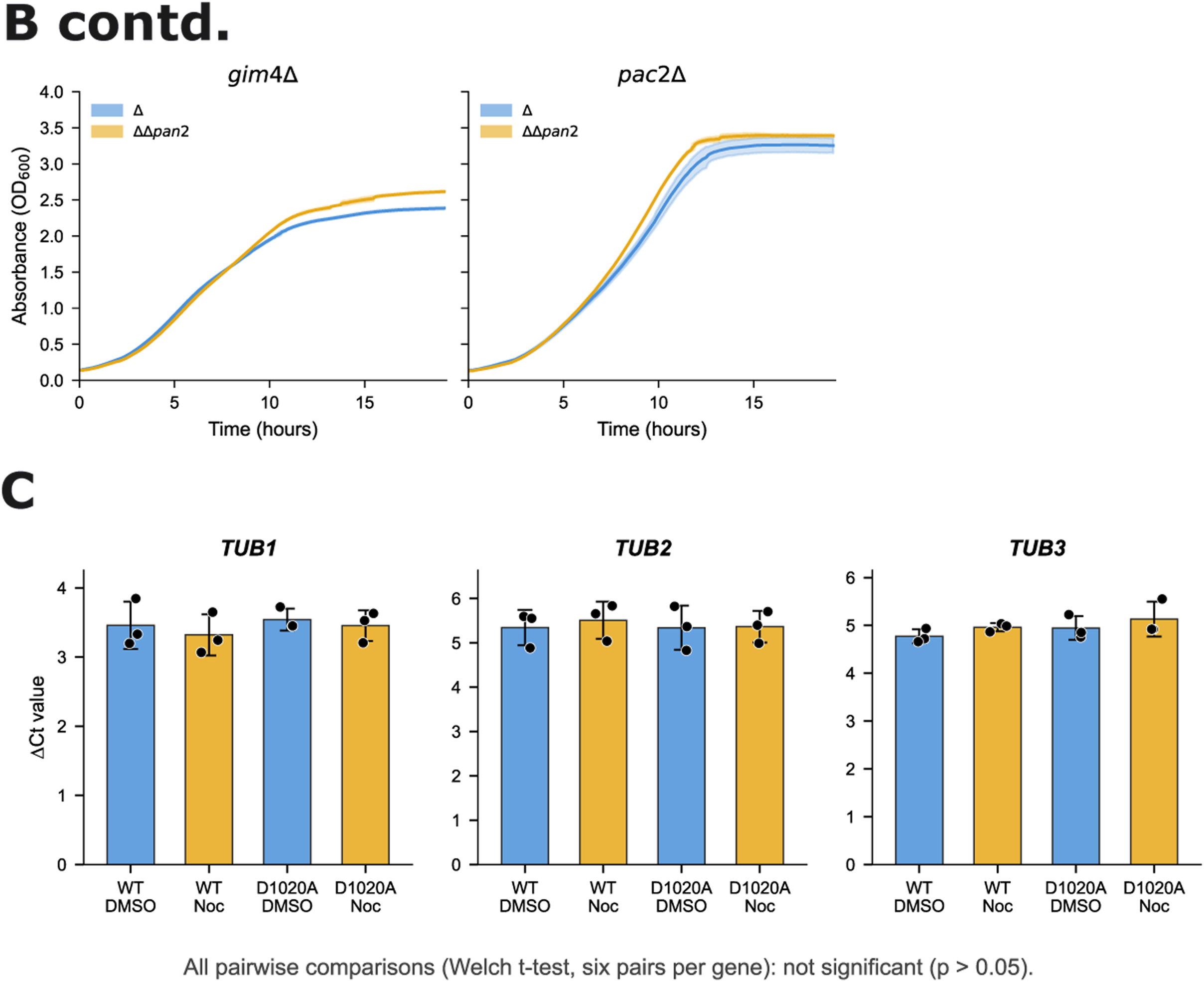
Genetic interaction analysis and tubulin mRNA levels. **(A)** CellMap *PAN2*/*PAN3* genetic-interaction analysis (Costanzo *et al*., 2016). Top: heatmap of *PAN2* (YGL094C) and *PAN3* (YKL025C) genetic interactions across the strongest interactors; darker blue = stronger aggravating, darker yellow = stronger alleviating. Bottom: Cytoscape functional-enrichment network of common interactors. The chromosome-segregation cluster includes *APC1*, *BUB3*, *CBF1*, *MPS1*, *SLI15*, *SMC3*, *TUB2*, and *TUB3*, with subcategories for spindle, kinetochore, microtubule cytoskeleton, mitotic spindle assembly checkpoint signaling, mitotic sister chromatid separation, polymeric cytoskeletal fiber, mitotic nuclear division, and metaphase/anaphase transition. Additional clusters implicate cytoskeletal protein binding (*ARP2*, *CIN1*, *GIM3*, *GIM5*, *PAC10*, *RBL2*, *YKE2*), RNA polymerase II-specific transcription factor binding (*CBF1*, *GCR2*, *SWI1*), and secretion (*ACT1*, *SEC1*, *SSO2*, *VPS27*). Edge thickness reflects statistical strength of association. **(B)** DMSO control for double mutants. Growth curves of single-mutant (blue) and *pan2Δ* double-mutant (orange) strains in SC media + DMSO at 30 °C, layout matching Figure 2B (WT vs *pan2Δ*; *tub3Δ*, *cin1Δ*, *cin2Δ*, *cin4Δ*, *gim3Δ*, *gim4Δ*, *pac2Δ*). Double mutants show no significant growth defect relative to single mutants in the absence of drug, confirming that the synthetic genetic interactions revealed in Figure 2B are nocodazole-stress-conditional. Absorbance (OD₆₀₀) plotted with shaded ± SD. Data: 6 technical replicates per experiment, 3 biological replicates. **(C)** qRT-PCR analysis of tubulin mRNA levels. ΔCt values for *TUB1*, *TUB2*, and *TUB3* in *pan2Δ* + Pan2-WT and *pan2Δ* + Pan2-D1020A cells after 6 h of 6 μM nocodazole or DMSO. No significant change in steady-state transcript abundance is observed between catalytic-active and catalytic-dead Pan2 conditions, arguing against catalytic-activity-dependent regulation of tubulin transcripts by PAN and implicating post-transcriptional mechanisms acting on other targets. Mean ± SD; all pairwise comparisons (two-tailed Welch’s t-test, six pairs per gene) not significant (p > 0.05). Data: 3 biological replicates with technical triplicates.

**Figure S3:**
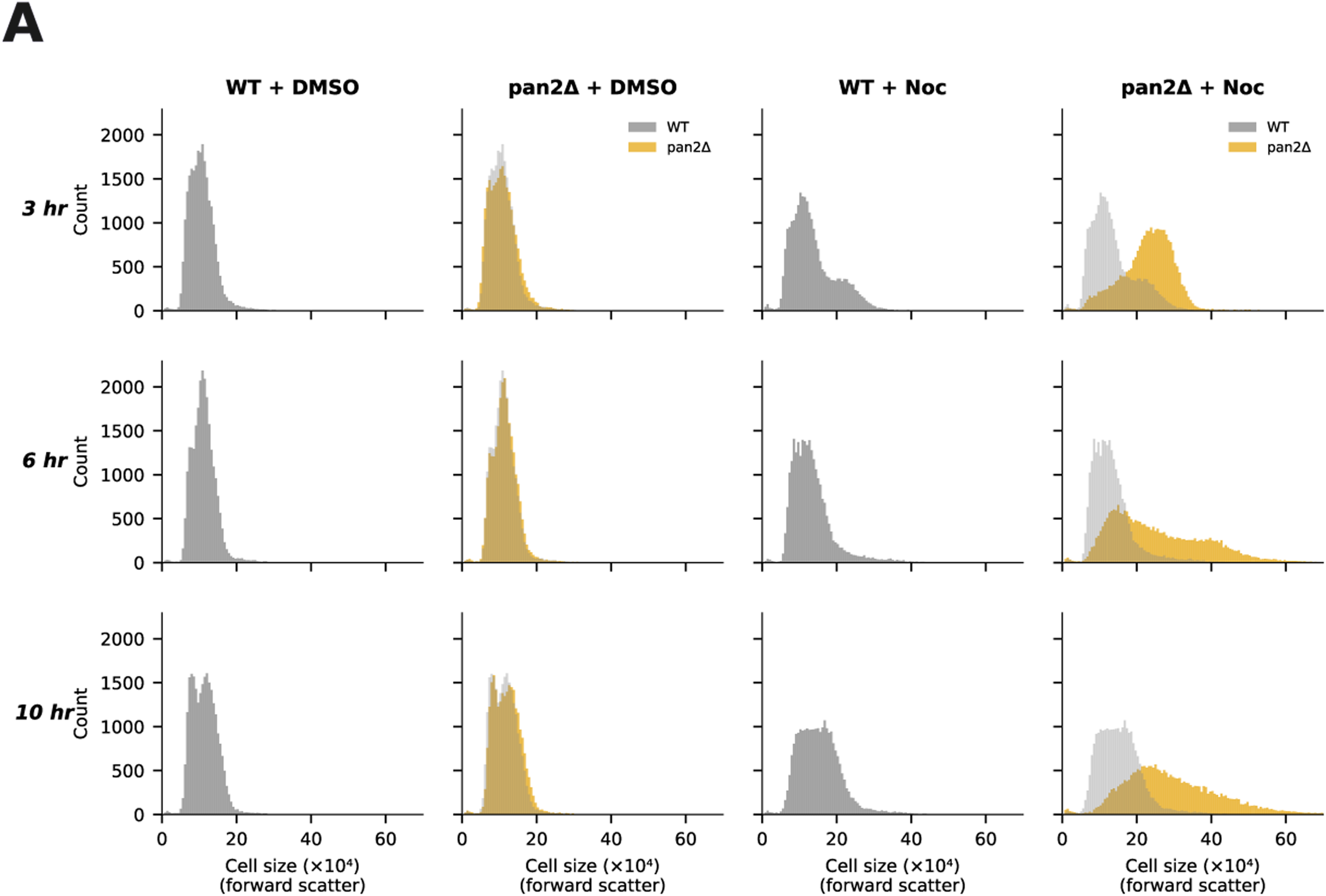
Forward-scatter cell-size analysis. Forward-scatter (FSC) intensity histograms of wild-type and *pan2Δ* cells cultured ± 6 μM nocodazole for 3, 6, and 10 hours. FSC correlates with cell volume; the x-axis is cell size (×10⁴, FSC-A) and the y-axis is cell count. Four conditions are shown across each time point: WT + DMSO, *pan2Δ* + DMSO, WT + Noc, *pan2Δ* + Noc. Gray WT overlays on *pan2Δ* + Noc histograms facilitate direct comparison. *pan2Δ* cells under nocodazole exhibit a progressive rightward shift (larger mean FSC) as early as 3 hours that becomes more pronounced at 6 and 10 hours, while DMSO-treated cells of both genotypes show no size difference. Data: representative histograms from 3 biological replicates; ≥10,000 cells per sample.

**Figure S4:**
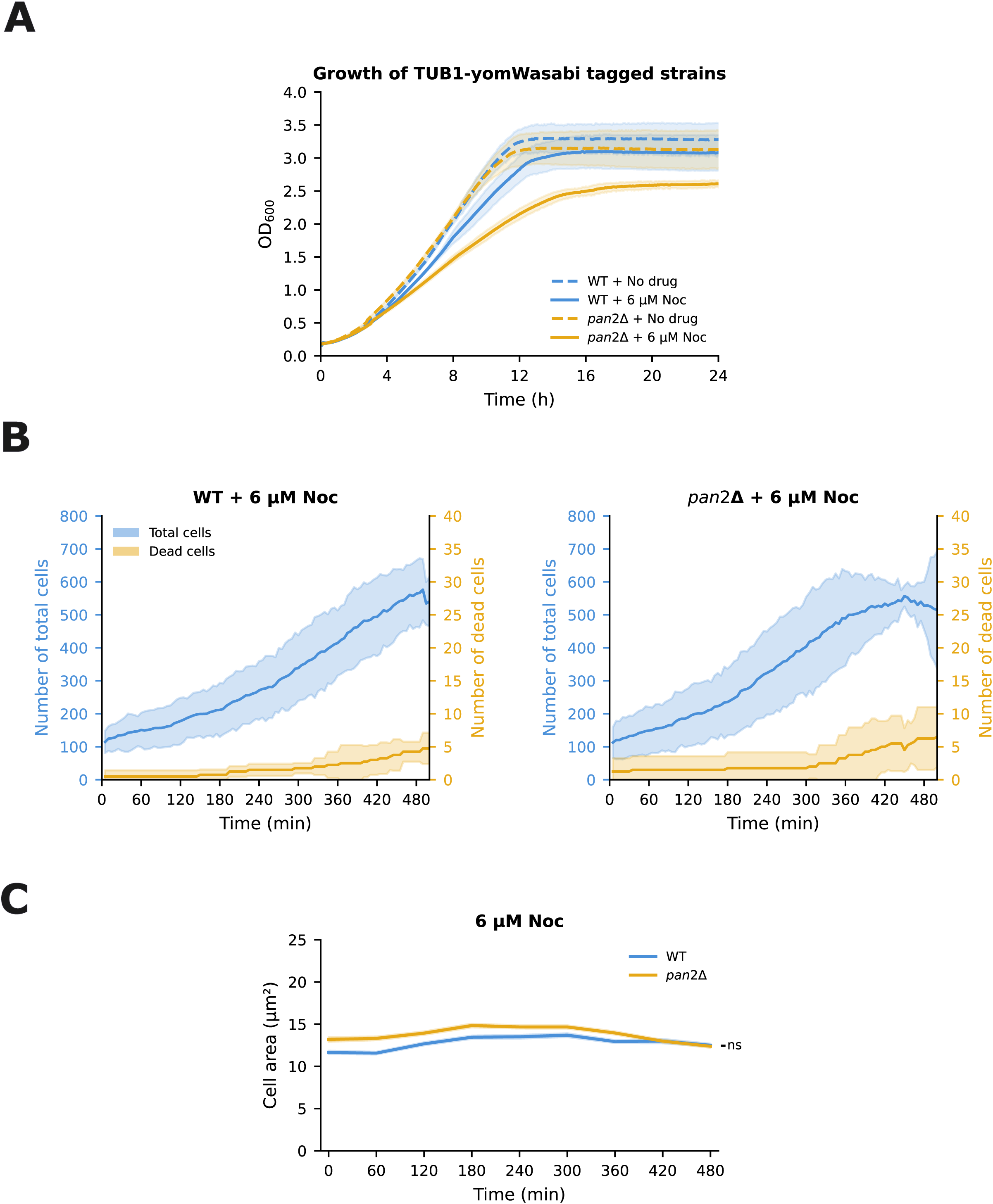
Live-cell imaging analysis of *TUB1*-yomWasabi-tagged strains. **(A)** Growth curves of *TUB1*-yomWasabi-tagged WT and *pan2Δ* strains in SC media + DMSO or 6 μM nocodazole over 24 h (OD₆₀₀). The tagged *pan2Δ* strain retains nocodazole sensitivity comparable to untagged *pan2Δ*; the tagged WT strain shows only a mild basal slow-growth phenotype, confirming that the fluorescent tag does not appreciably alter the drug-sensitivity phenotype. Mean ± SD shaded. Data: 6 technical replicates per experiment, 3 biological replicates. **(B)** Live-cell imaging quantification at 6 μM nocodazole. Total cell count (left y-axis, blue) and dead cell count (right y-axis, orange) over 480 minutes for WT (left) and *pan2Δ* (right). Under static (non-aerated) imaging conditions, 6 μM nocodazole did not produce a significant difference in cell number or dead cells between WT and *pan2Δ*, in contrast to the liquid-shaking growth assay. We attribute this to reduced effective drug exposure under static conditions, motivating the increased dose (12 μM) used in Figure 4. Mean ± 95% CI shaded. Data: ≥10 fields of view per condition; 4 biological replicates. **(C)** Mean cell area (μm²) over time at 6 μM nocodazole for WT (blue) and *pan2Δ* (orange). No significant differences (n.s.) at this lower drug concentration under static imaging, in contrast to the robust phenotype at 12 μM nocodazole (Figure 4C). Mean ± SEM shaded. Data: n > 100 cells per condition per time point; 3 biological replicates.

**Figure S5:**
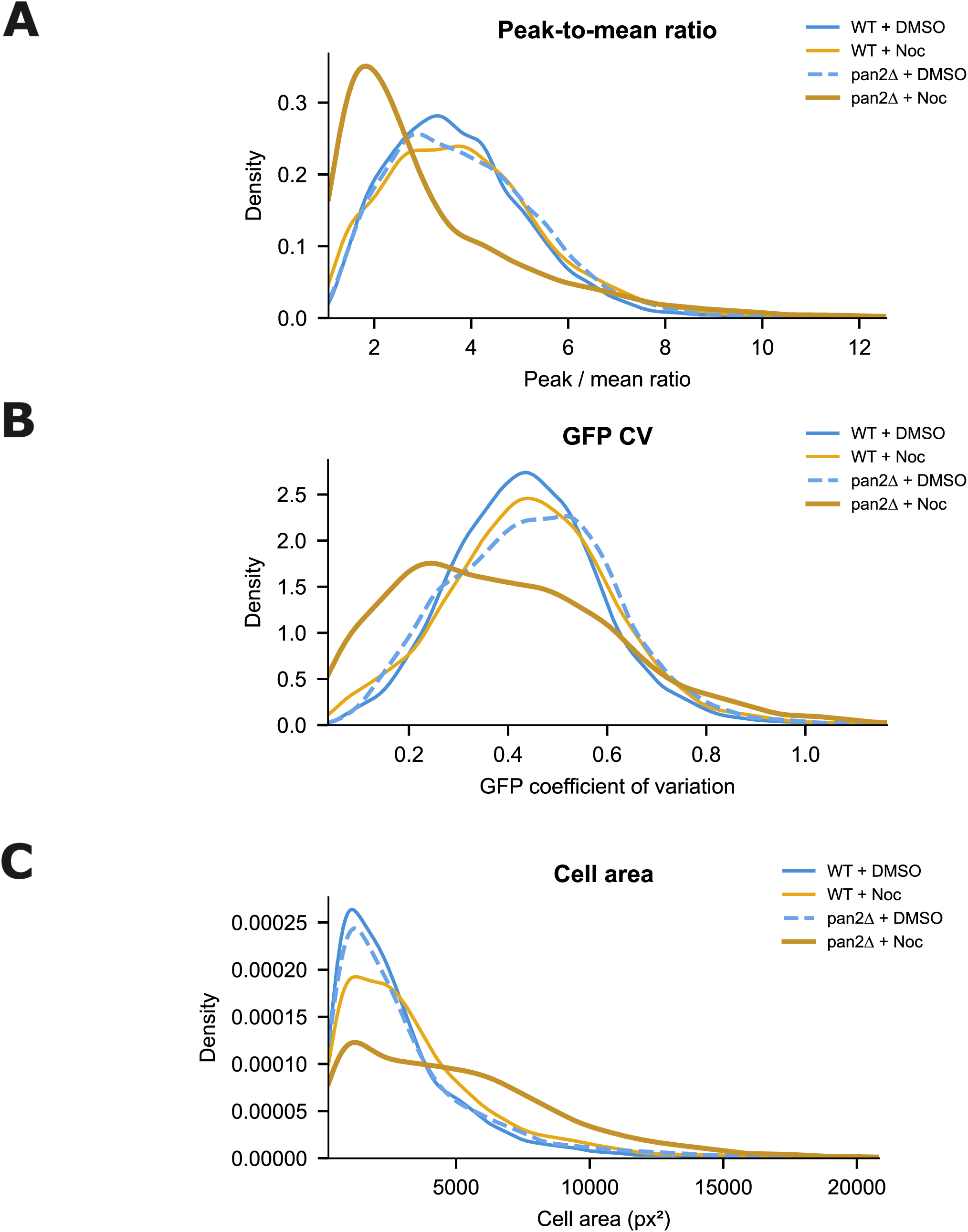
Kernel density estimation of microtubule organization metrics. Related to Figure 5. **(A)** Kernel density estimation (KDE) of peak-to-mean GFP intensity ratios across the four imaging conditions (WT + DMSO, WT + Noc, *pan2Δ* + DMSO, *pan2Δ* + Noc). *pan2Δ* + Noc cells show a leftward-shifted, narrow distribution centered at low ratios (∼2), consistent with loss of concentrated GFP-tubulin puncta, whereas WT + Noc cells maintain a broader distribution with substantial fractions at high ratios (∼3–5). **(B)** KDE of GFP coefficient of variation (CV). *pan2Δ* + Noc cells show a narrower distribution shifted toward lower CV values, consistent with more uniform (diffuse) GFP-tubulin fluorescence and loss of discrete microtubule structures. WT + Noc cells maintain higher CV values reflecting organized punctate signal. **(C)** KDE of cell area (px²). *pan2Δ* + Noc cells exhibit a rightward-shifted distribution (larger cells) consistent with prolonged G2/M arrest. WT + DMSO and *pan2Δ* + DMSO distributions overlap, indicating that cell enlargement is drug-dependent. Data for all panels: n’s as in Figure 5; pooled from 3 independent biological replicates with dead cells excluded.

**Figure S6:**
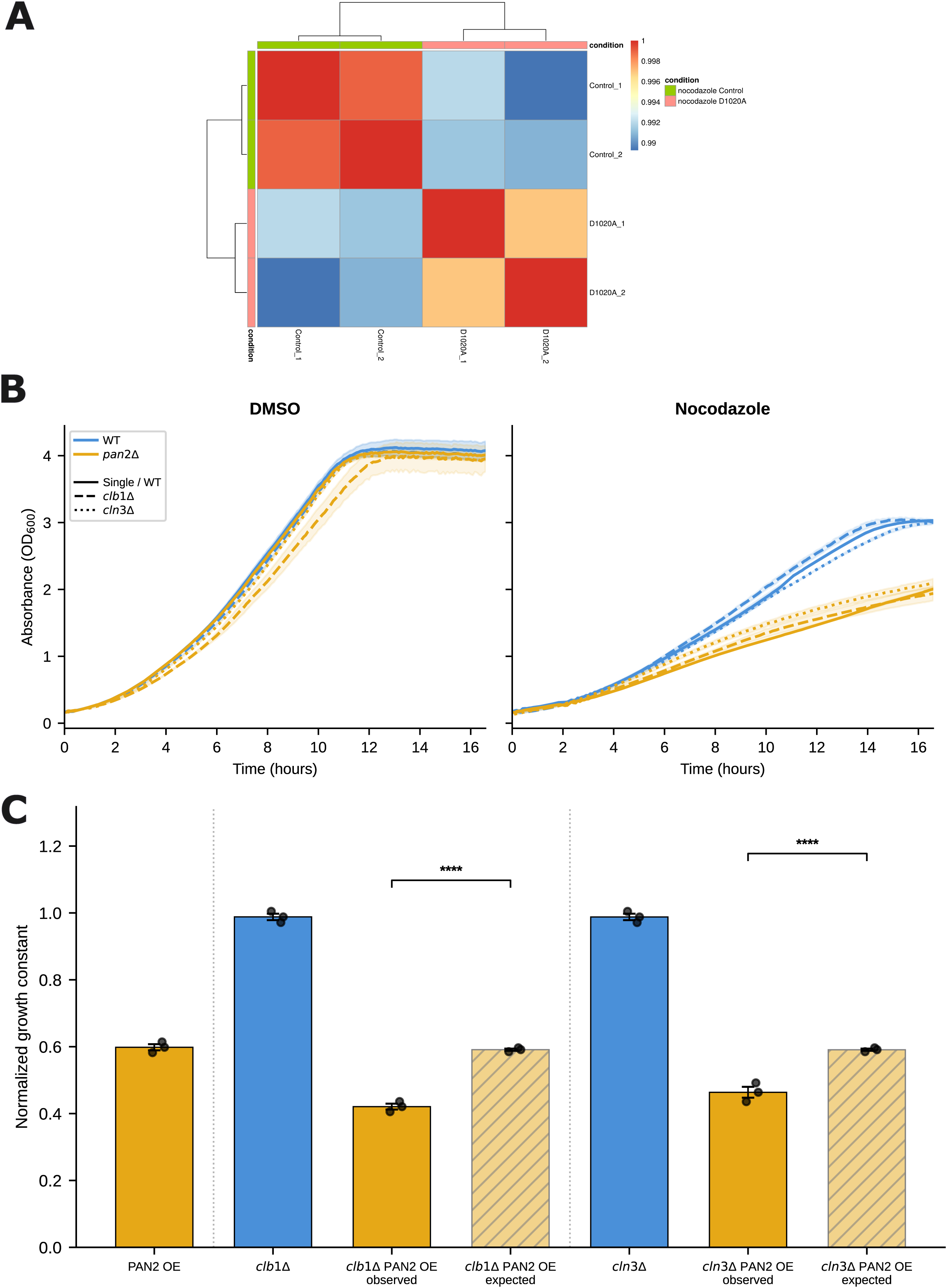
RNA-seq quality control and cyclin genetic interactions. Related to Figure 6. **(A)** Spearman correlation matrix of RNA-seq replicates from *pan2Δ* + Pan2-WT (Control_1, Control_2) and *pan2Δ* + Pan2-D1020A (D1020A_1, D1020A_2) samples treated with 6 μM nocodazole for 6 h. High intra-genotype correlation confirms reproducibility of expression measurements across replicates. **(B)** Growth curves of cyclin deletion mutants ± *pan2Δ* in SC media + DMSO (left) and SC + 6 μM nocodazole (right). Strains: WT (blue solid), *pan2Δ* (orange solid), *clb1Δ* (blue dashed), *cln3Δ* (blue dotted), and their *pan2Δ* double mutants (orange dashed/dotted). Growth of each double mutant matches the *pan2Δ* single mutant in both DMSO and nocodazole, indicating that loss of either cyclin alone does not modify *pan2Δ* sensitivity, motivating the *PAN2*-overexpression approach in Figure 6D. Absorbance (OD₆₀₀) plotted with shaded mean ± SEM. Data: 6 technical replicates per experiment, 3 biological replicates. **(C)** Quantitative epistasis analysis of cyclin deletions with *PAN2* overexpression. Normalized growth constants (in nocodazole) for *PAN2* OE alone, *clb1Δ* alone, and *clb1Δ* + *PAN2* OE (observed vs. expected from a multiplicative null); same comparisons for *cln3Δ*. Observed double-mutant growth is significantly lower than the multiplicative-null expectation for both cyclin combinations, confirming aggravating (synergistic) genetic interactions. Mean ± SD; ****p < 0.0001 by two-tailed Student’s t-test (observed vs. expected). Data: 6 technical replicates per experiment, 3 biological replicates.

