## Supplementary material for "PAN2-PAN3 deadenylase activity is selectively required for mitotic robustness under microtubule stress": Yeast-codon-optimized human PAN2 sequence (FASTA).

Sequence of human PAN2 (HsPAN2) codon-optimized for expression in

Saccharomyces cerevisiae. Synthesized by Twist Biosciences and cloned

into Gateway destination vectors as described in Methods. Encodes the

full-length 1202-amino-acid human PAN2 protein.

>HsPAN2_codon_optimized_yeast

ATGAACTTCGAGGGATTGGATCCAGGATTAGCGGAGTATGCACCCGCGATGCATTCTGCG

CTAGACCCCGTGTTAGATGCACACTTAAACCCCTCTCTTCTGCAGAATGTAGAATTGGAC

CCCGAGGGCGTAGCATTAGAGGCTCTGCCGGTTCAGGAGTCCGTACACATAATGGAAGGG

GTGTACTCGGAGTTACACTCAGTTGTGGCTGAGGTAGGCGTACCTGTGTCCGTTTCACAC

TTCGACTTACACGAGGAGATGCTGTGGGTAGGCAGTCACGGCGGCCATGCCACATCATTC

TTCGGACCTGCCCTTGAGAGGTATTCAAGCTTCCAAGTAAATGGCAGTGACGATATTCGT

CAGATACAGTCCCTAGAGAATGGGATACTATTCCTTACAAAGAACAACTTAAAGTATATG

GCACGTGGCGGCCTGATAATCTTTGACTACTTGTTGGACGAGAACGAAGACATGCATTCG

TTGCTTTTGACAGACAGCTCCACACTATTAGTCGGCGGATTGCAGAACCATATCATAGAG

ATTGACTTGAACACTGTACAGGAAACGCAGAAGTATGCGGTCGAGACCCCTGGTGTGACC

ATAATGAGGCAGACAAACAGGTTCTTCTTCTGCGGACACACAAGTGGTAAAGTAAGCCTG

AGAGACCTTAGAACTTTCAAAGTCGAGCATGAGTTCGACGCATTTTCGGGTAGCCTAAGT

GACTTCGACGTACATGGCAACCTACTTGCTGCATGCGGATTTTCTAGCAGGTTGACTGGG

TTAGCTTGTGACAGGTTCTTAAAGGTCTACGACCTGAGGATGATGCGTGCAATTACTCCG

CTACAGGTTCACGTAGACCCCGCCTTCCTTAGGTTCATACCTACATACACGTCAAGACTG

GCAATAATATCCCAGTCAGGCCAGTGCCAGTTCTGCGAGCCAACCGGACTGGCCAATCCG

GCAGACATTTTCCACGTGAACCCTGTCGGACCTCTTCTTATGACGTTCGACGTCAGTGCA

TCGAAACAGGCACTAGCTTTCGGCGATTCTGAGGGCTGCGTACATTTATGGACTGACAGC

CCTGAGCCATCGTTCAACCCATACTCCAGGGAGACCGAGTTCGCATTGCCTTGTTTAGTA

GACTCCTTGCCACCTCTAGACTGGTCGCAGGACCTTTTACCTCTAAGTTTAATACCTGTA

CCCTTGACTACCGACACCTTGTTAAGTGACTGGCCAGCTGCGAACTCCGCACCAGCTCCA

CGTAGGGCTCCACCGGTTGACGCCGAGATTCTGCGTACAATGAAGAAGGTGGGCTTTATT

GGCTATGCCCCTAATCCGCGTACACGTCTGAGGAACCAGATCCCTTATCGTCTAAAAGAG

TCCGACAGTGAGTTCGACTCATTCTCCCAGGTAACCGAGTCCCCGGTTGGTCGTGAAGAG

GAGCCTCACTTACATATGGTCTCCAAGAAATACAGGAAGGTTACAATTAAATACAGTAAA

TTGGGTCTAGAGGACTTCGACTTCAAGCACTACAACAAGACTTTATTTGCGGGACTGGAG

CCACATATACCTAACGCCTACTGCAATTGCATGATACAGGTTCTATATTTTCTAGAGCCC

GTACGTTGTCTGATACAGAATCACCTGTGCCAGAAAGAGTTTTGCTTAGCTTGCGAGCTA

GGCTTTCTATTTCACATGTTGGACTTGTCACGTGGAGACCCATGCCAGGGCAATAACTTC

TTGAGGGCATTCAGGACTATACCCGAGGCCTCAGCCTTAGGCTTGATACTGGCTGACAGC

GATGAGGCATCGGGGAAGGGAAATTTGGCCCGTTTAATACAGCGTTGGAATCGTTTTATC

CTTACTCAGCTGCACCAGGACATGCAGGAGTTGGAAATCCCTCAGGCCTACAGGGGAGCT

GGTGGGTCAAGCTTCTGCAGCTCTGGAGACTCTGTAATAGGCCAGTTGTTTTCATGCGAG

ATGGAGAACTGCTCTCTATGCAGGTGCGGCTCCGAAACGGTAAGGGCGTCCTCTACACTT

CTATTCACCTTATCATACCCTGATGGAAGTAAGTCTGATAAAACCGGAAAGAACTACGAC

TTCGCACAGGTGTTAAAGAGGTCGATCTGCTTGGACCAGAACACCCAAGCTTGGTGCGAC

ACGTGCGAGAAATACCAGCCCACTATACAGACACGTAATATCAGACACCTTCCGGACATC

TTAGTCATTAATTGCGAGGTAAACTCTAGCAAAGAGGCCGACTTCTGGAGAATGCAGGCC

GAGGTAGCGTTCAAGATGGCAGTGAAGAAGCACGGTGGAGAGATAAGTAAGAACAAGGAG

TTTGCCCTTGCGGACTGGAAAGAGCTAGGGAGTCCAGAGGGCGTACTAGTCTGTCCATCC

ATCGAGGAGCTTAAGAATGTCTGGTTGCCGTTTTCCATTAGGATGAAGATGACGAAGAAC

AAAGGGCTAGACGTCTGTAATTGGACAGACGGCGACGAGATGCAGTGGGGTCCAGCTCGT

GCCGAGGAAGAGCACGGAGTTTATGTGTACGACCTAATGGCAACCGTCGTGCACATACTT

GACAGCCGTACAGGCGGGAGTCTAGTGGCGCACATAAAAGTAGGAGAGACCTACCACCAA

AGGAAAGAAGGAGTAACGCACCAACAGTGGTATCTTTTCAATGACTTTCTAATTGAGCCT

ATCGACAAACATGAGGCAGTACAGTTCGATATGAATTGGAAGGTACCAGCCATCTTATAT

TACGTAAAGAGGAACCTAAACAGTAGGTACAATTTAAACATAAAGAACCCCATCGAGGCC

TCTGTTCTTTTAGCGGAAGCGTCGTTAGCCCGTAAGCAGAGGAAGACACACACAACTTTT

ATTCCCTTAATGCTGAACGAGATGCCACAGATCGGGGATTTGGTCGGATTGGACGCAGAG

TTCGTAACCCTGAACGAGGAAGAGGCTGAGTTGAGGAGTGACGGCACCAAGTCCACGATT

AAGCCTTCACAGATGTCAGTAGCAAGGATAACGTGCGTCCGTGGTCAGGGACCGAATGAG

GGCATCCCTTTCATCGATGACTACATATCTACCCAGGAGCAGGTCGTTGACTATCTGACT

CAGTATTCTGGGATCAAGCCAGGCGACTTGGACGCTAAAATCAGCAGTAAGCACCTGACC

ACGTTAAAGTCAACCTATCTAAAGCTAAGATTCTTAATAGACATCGGAGTCAAGTTCGTC

GGTCACGGGCTACAGAAGGACTTCAGGGTCATTAACCTTATGGTCCCGAAGGACCAGGTC

TTGGACACAGTGTACCTTTTCCACATGCCAAGGAAGAGAATGATATCCTTAAGGTTCTTA

GCCTGGTACTTCTTAGACCTTAAAATACAGGGTGAGACCCACGACTCCATAGAGGACGCT

CGTACGGCGTTGCAGTTATACAGGAAATATCTAGAGCTGTCAAAGAACGGCACTGAGCCA

GAGAGTTTTCACAAGGTGTTAAAAGGCCTTTACGAGAAAGGAAGGAAAATGGACTGGAAG

GTGCCAGAACCCGAGGGGCAGACTAGCCCGAAGAATGCAGCCGTGTTCTCTAGTGTGTTA

GCCTTATAG
